# Ciliated protozoa from industrial WWTP activated sludge: a biodiversity survey including trophic interactions and redescription of *Bakuella subtropica* (Spirotrichea, Hypotrichia) according to Next Generation Taxonomy

**DOI:** 10.1101/2021.06.01.446513

**Authors:** Wanying Liao, Valentina Serra, Leandro Gammuto, Francesco Spennati, Gualtiero Mori, Giulio Munz, Letizia Modeo, Giulio Petroni

**Author notes:** Correspondence to Giulio Petroni) or etizia Modeo.

## Abstract

Optimization of wastewater treatment with biological processes is a fundamental challenge of modern society. During past years new technologies have been developed for the purpose and prokaryotic organisms involved in the process extensively investigated. Nevertheless, relatively few studies so far analysed the protozoan community in these systems using modern integrative approaches, despite its obvious role in shaping ecological dynamics and, possibly, process efficiency. In the present study, we characterized the ciliate community in biological reactors of an Italian industrial (tannery) wastewater treatment plant (WWTP) applying modified Ludzack-Ettinger (MLE) process. This plant is characterized by moderate salinity, high solids retention time and high concentration of organic compounds, including a significant recalcitrant fraction. We performed the morphological and 18S rDNA characterizations of almost all the 21 ciliates retrieved along a one-year sampling period, and provided preliminary data on species occurrence, community dynamics, and trophic interactions. Only 16 species were observed on the sample collection day and most of them had an occurrence higher than 50%. The most frequently occurring and highly abundant organisms were *Aspidisca* cf. *cicada*, *Euplotes* spp., *Paramecium calkinsi*, and *Phialina* sp. *Cyclidium* cf. *marinum* was only found on a single date and its presence was possibly related to a summer break-induced perturbation. All the species showed the capability to survive the short oxic/anoxic cycling typical of the studied WWTP process. Intriguingly, some of them (i.e., *Bakuella subtropica* and *Trochiliopsis australis*) turned out to be species isolated from brackish natural environment rich in organic load as well. As for *B. subtropica*, we provided an emended redescription according to the most recent taxonomy standards that include also mitogenomic sequencing.

## Introduction

As one major member of the microorganism community in the activated sludge system used for biological wastewater treatment, ciliated protozoa play important roles and have been extensively studied under this respect (e.g., Curds and Cockburn, 1970; Curds, 1973a, b; Esteban et al., 1991; Madoni et al., 1993; Salvadó and Gracia, 1993; Madoni, 1994; Salvadó, 1994; Martín-Cereceda et al., 1996; Lee et al., 2004; Liu et al., 2008; Pérez-Uz et al., 2010; Dubber and Gray, 2011; Madoni, 2011; dos Santos et al., 2014; Foissner, 2016). On the contrary, the taxonomy of activated sludge ciliates has generally received little attention. Indeed, most works are still frequently based on microscopic examination of biomass, which is a fast, simple, convenient but low-precision method, although integrative taxonomy (i.e., the multimethod taxonomy performed through morphological-ultrastructural study combined with phylogenetic analysis based on molecular markers) has been widely recognized as the standard approach for species identification (Foissner, 2016). For this reason, it is easy to overlook the undescribed species in wastewater treatment plants (WWTPs), and some poorly known species still lack redescriptions based on modern techniques and criteria (Aescht and Foissner, 1992; Leitner and Foissner, 1997; Guggiari and Peck, 2008; da Silva Paiva et al., 2016; Foissner, 2016).

The taxonomic composition and population distribution of protozoa are directly related to the type of wastewater treatment process applied, the operating conditions, and the composition of the wastewater treated (Curds, 1973a, b; Madoni et al., 1993; Madoni, 1994, 2011; Foissner, 2016). In this context, a widely used application is the sludge biotic index proposed by Madoni (1994). It is an index based on the structure and abundance of the microfauna inhabiting the activated sludge, and it provides important information for monitoring activated sludge plants performance. One advantage of this approach is that it generally does not need a precise taxonomic identification at species level of the involved organisms, so a fast *in vivo* check of the sample is sufficient. In Madoni’s work, ciliates in activated sludge have been subdivided into four groups on the basis of their behaviour (feeding and movement habits), that is (1) free-swimming bacterivores; (2) crawling bacterivores; (3) sessile bacterivores; and (4) carnivores involving both free-swimming (such as members of the genera *Amphileptus*, *Litonotus*, and *Trachelophyllum*) and sessile ones represented by suctorian ciliates like *Acineta* and *Podophrya*. However, many omnivorous ciliates feeding both on bacteria and other bacterivores have been ignored in Madoni’s classification. Because of their unique position in the food web, these organisms should be separately considered, even if the selective food preferences of these organism remain unclear. In the present work, we tried to provide a more refined and reliable behavioural repartition of the ciliate community in the activated sludge system, as required to better understand the interdependencies among these organisms and within the microbial community.

In addition, so far, many studies have only focused on ciliated protozoa in the conventional activated sludge process, which includes only one biological oxidation tank for the reduction of the organic matter present in the wastewater (e.g., Curds, 1973a; Esteban et al., 1991; Madoni et al., 1993; Salvadó and Gracia, 1993; Madoni, 1994; Salvadó, 1994; Martín-Cereceda et al., 1996). Few studies have evaluated the activated sludge microfauna in WWTPs that use modified Ludzack-Ettinger (MLE) process for removal of organic matter, ammonia, and nitrate/nitrite through the combined anoxic-aerobic zones, and few studies are dealing with industrial wastewater. However, the studies (Liu et al., 2008; Pérez-Uz et al., 2010; Dubber and Gray, 2011) on WWTPs applying MLE process revealed that a significantly different protozoan community could be observed with the introduction of anoxic stage, and some biological indicators of ciliates used in conventional activated sludge plants could not be directly applied. Also, the toxic substances present in the industrial wastewater seem to challenge previous conclusions drawn from investigations on domestic wastewater (Papadimitriou et al., 2007; dos Santos et al., 2014). Moreover, unfortunately, all current studies on modified conventional activated sludge system only analyse samples collected from the aeration tank; and, to the best of our knowledge, no attempts have been made to study the difference in the ciliate community shaped by the anoxia/anaerobiosis effect, although this difference might also be insignificant due to water recirculation.

Tannery wastewaters, regardless of the type of industrial process (chromium or vegetable), are among the most difficult to treat, basically due to their recalcitrance and/or their toxicity towards the microfauna (Lofrano et al., 2013). Different types of wastewater provide different microorganisms. Fungi and microbial community have already been studied for WWTPs treating tannery wastewaters (Giordano et al., 2016; Tigini et al., 2018), while there is less knowledge on the ciliate community.

The main goal of the present study was to investigate the composition of ciliate community in an industrial WWTP applying MLE process to treat tannery wastewater and to provide preliminary insights on population dynamics and trophic interactions. We followed integrative taxonomy for species identification in order to provide an unambiguous checklist of the species present in this artificial habitat. Moreover, an attempt was also made for understanding the influence of oxygen concentration level on ciliate community in the specific WWTP condition by separately collecting samples from the nitrification tank (which is regarded as an oxic environment) and the denitrification tank (which is considered to be an anoxic environment). Finally, among the various retrieved ciliate species, *Bakuella subtropica* was redescribed based on the recently proposed taxonomic standard approach, i.e., the next generation taxonomy (NGTax) (Serra et al., 2020), and a critical revision of the genus was conducted, in line with previous papers from our group (e.g., Serra et al., 2021a; Serra et al., 2021b),with a proposal of some taxa synonymization.

## Results

As predictable from the high recirculation between nitrification and denitrification tanks, the ciliate community structure, at the same sampling times, was similar in the two tanks (Table S1). We consequently considered the samplings as independent replicates of the same environment and merged the data into a single table (Table 1). Considering that ciliate presence was recorded using four relative abundance scores (0 to 3), any arithmetic average could not be performed. When the recorded scores of two replicates were inconsistent, the higher abundance class was reported in Table 1. Indeed, the few differences between two replicates were typically related to some rare organisms or to those difficult to detect (Table S1); thus, we considered more reliable to refer to the higher value between the two.

**Table 1.**
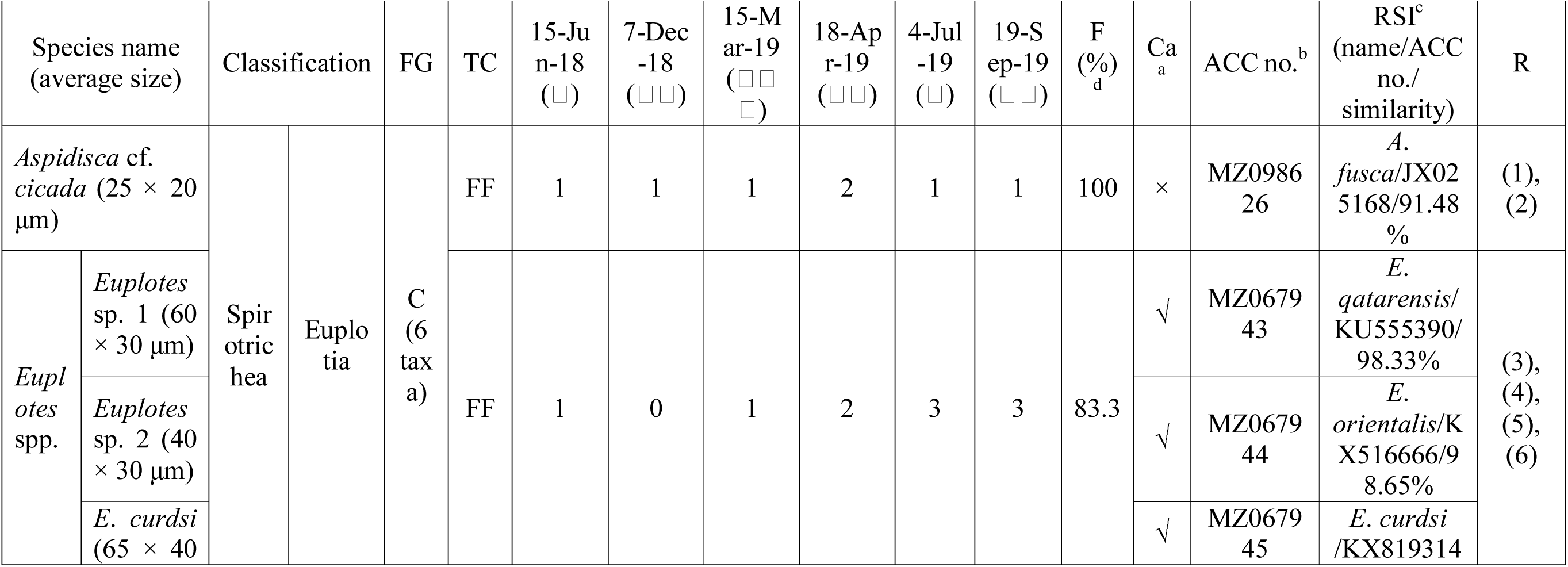
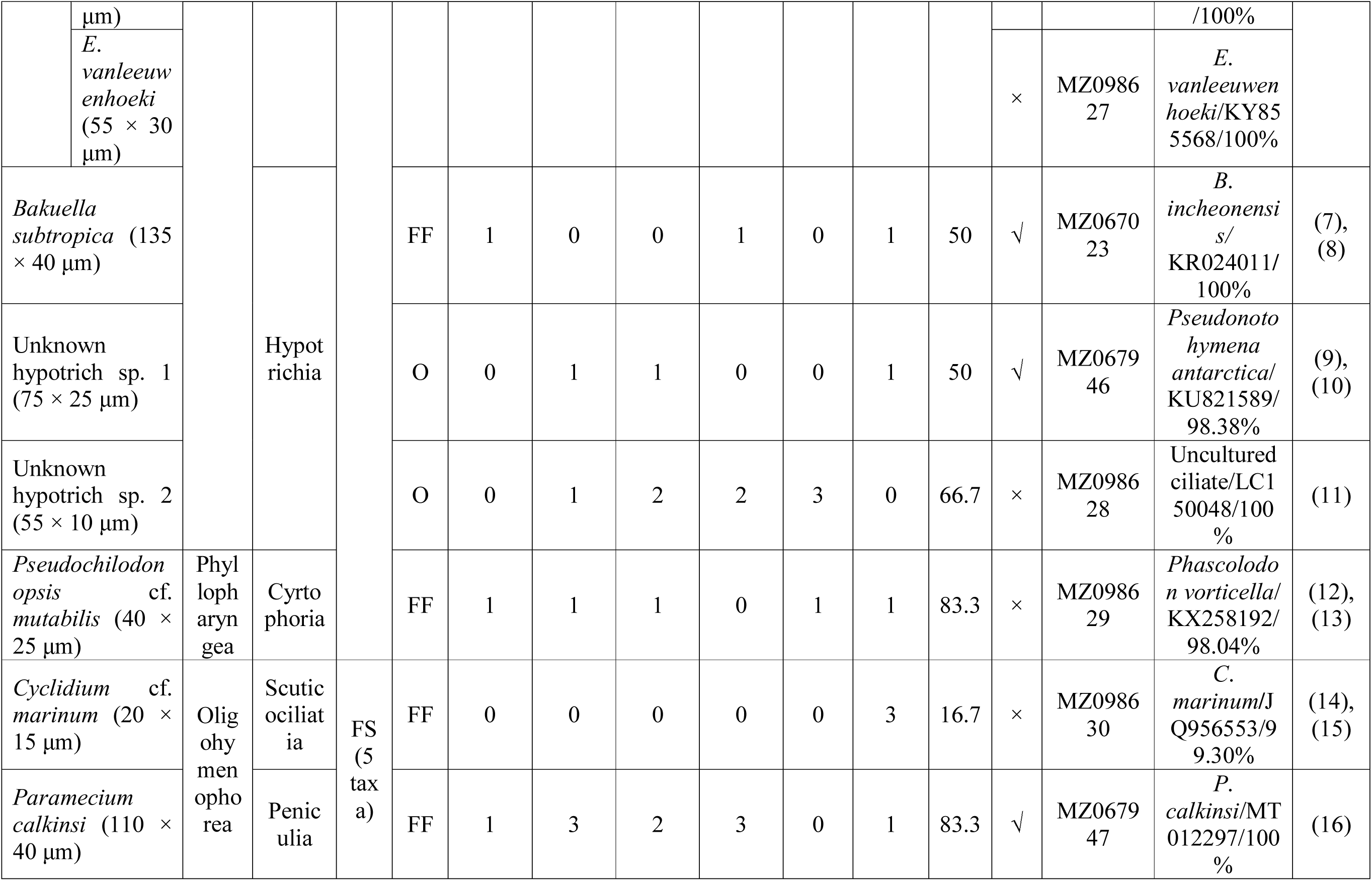
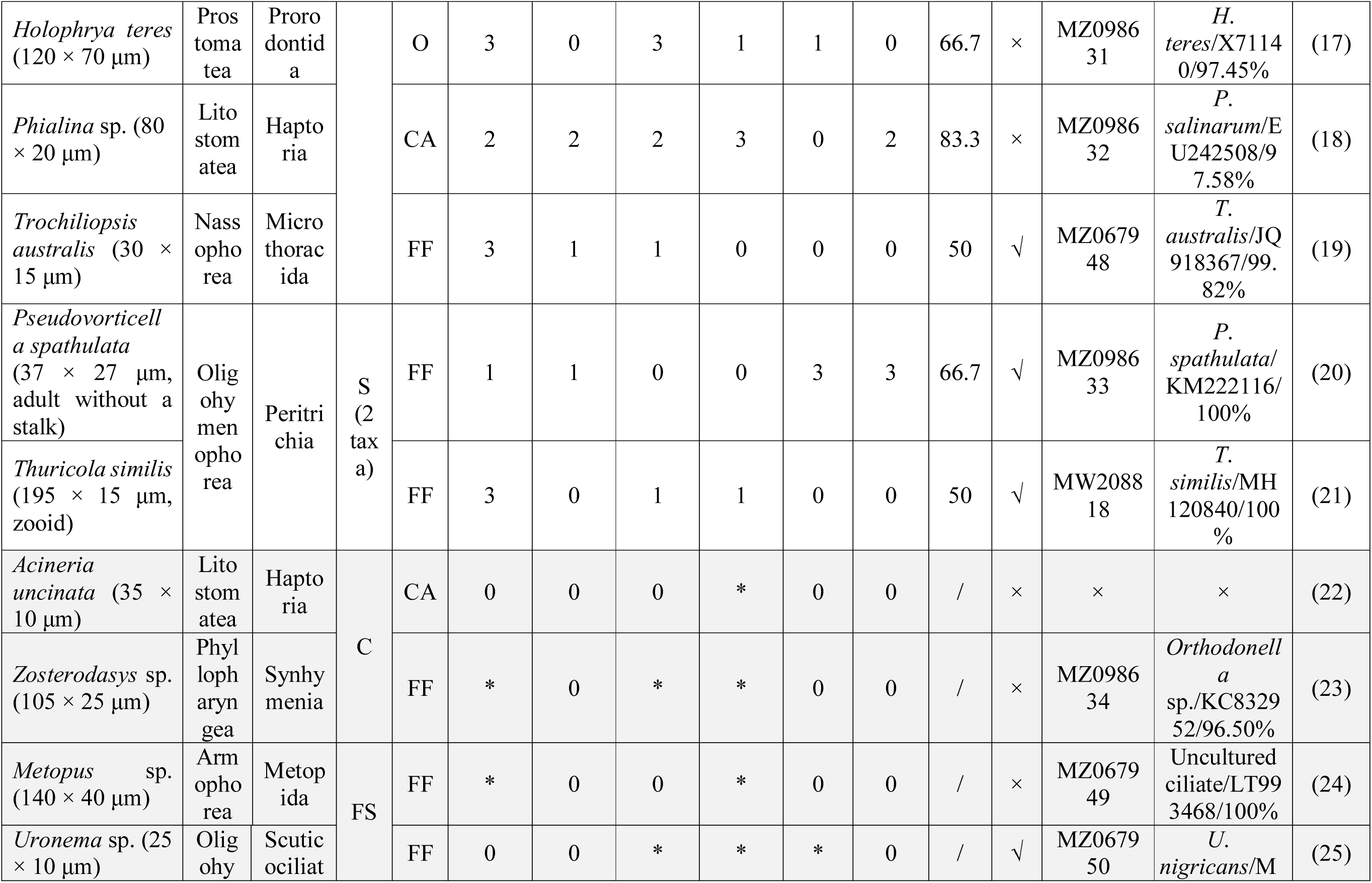
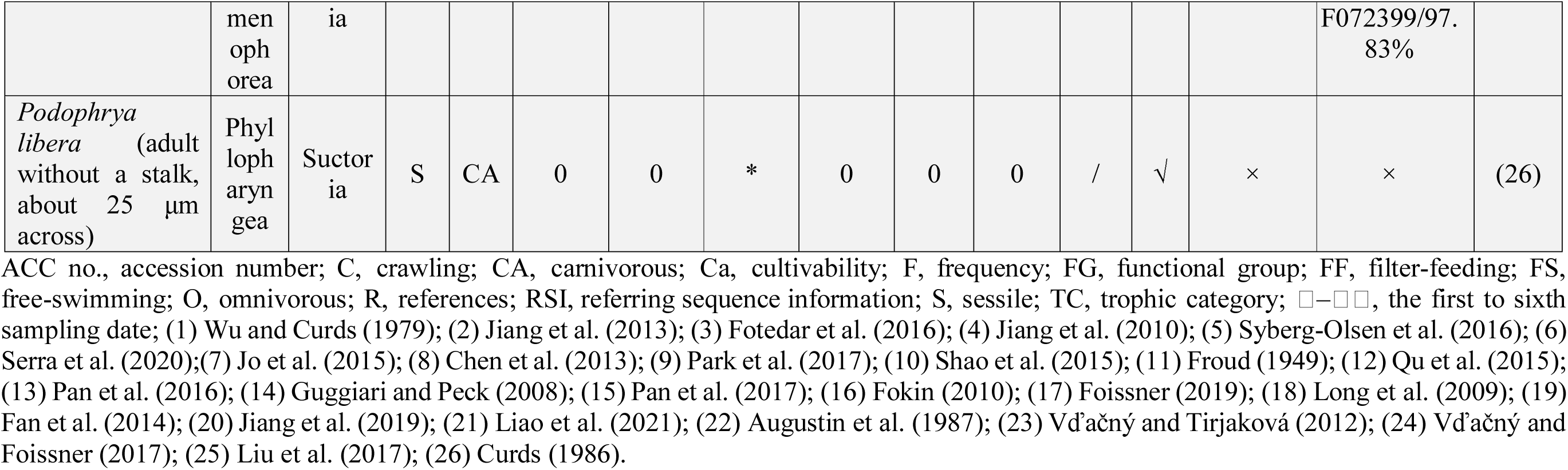
List of the species observed in Cuoiodepur WWTP with their relative abundance scores at the different sampling days. The species collected on the sampling day are displayed in the upper part of the table; then the species appeared only later during lab maintenance (shadowed in grey) are listed; the species were grouped firstly according to functional group, then to family membership, and, finally, to alphabetical order. The relative abundance scores of the species observed are presented as follows: 0 = absent in three subsamples on the sampling day; 1= present in few numbers (i.e., 1 to 9 cells could be collected from three subsamples within 30 min); 2 = present in moderate numbers (i.e., 10 to 29 cells could be collected within 30 min); 3 = present in massive numbers (i.e., more than 30 cells could be collected within 30 min). Asterisks indicate the original samples where the species of interest appeared after laboratory enrichment.√, succeeded; ×, failed; /, invalid data; a: monoclonal and/or polyclonal cultures could be established and maintained for three months in lab; b, GenBank accession numbers of newly obtained 18S rRNA gene sequences in the present study; c, the most similar sequence according to BLAST searching with priority always given to the taxonomically identified one (except LC150048 and LT993468, which are uncultured ciliates, but identical to Unknown hypotrich sp. 2 and *Metopus* sp., respectively). d, species frequency indicated as the occurrence rate (in percentage) of each species at the six sampling times. Note that, since the observed species of *Euplotes* were not discriminable *in vivo* due to their high morphological similarity, they were grouped as a single taxonomical unit (*Euplotes* spp.). In detail, the four *Euplotes* observed species, i.e., *Euplotes* sp. 1, *Euplotes* sp. 2, *E*. *curdsi*, and *E*. *vanleeuwenhoeki*, were observed throughout the study, with their first record on 15-Jun-2018, 15-Jun-2018, 4-Jul-2019, and 19-Sep-2019, respectively.

### Identity of the ciliate species found in Cuoiodepur WWTP

In total, 21 taxa were found in the biological process of Cuoiodepur WWTP and 18S rDNA molecular data were provided for 19 of them (Table 1). Except two non-identified hypotrichs, the remaining 19 taxa were identified at least to genus level, with 13 out of 19 identified to species level, i.e., *Aspidisca* cf. *cicada* (Spirotrichea, Euplotia), *Acineria uncinata* (Litostomatea, Haptoria), *Bakuella subtropica* (Spirotrichea, Hypotrichia), *Cyclidium* cf. *marinum* (Oligohymenophorea, Scuticociliatia), *Euplotes curdsi* (Spirotrichea, Euplotia), *Euplotes vanleeuwenhoeki*, *Holophrya teres* (Prostomatea, Prorodontida), *Paramecium calkinsi* (Oligohymenophorea, Peniculia), *Podophrya libera* (Phyllopharyngea, Suctoria), *Pseudochilodonopsis* cf. *mutabilis* (Phyllopharyngea, Cyrtophoria), *Pseudovorticella spathulata* (Oligohymenophorea, Peritrichia), *Thuricola similis* (Oligohymenophorea, Peritrichia), and *Trochiliopsis australis* (Nassophorea, Microthoracida). Among them, *Cyclidium* cf. *marinum*, *E*. *curdsi*, *E*. *vanleeuwenhoeki*, *P*. *calkinsi*, *P*. *spathulata*, and *T*. *australis* were undoubtedly identified solely based on their 18S rRNA gene sequences: four species showed a 18S rRNA gene sequence identical to their conspecifics’ sequences available in the database, and two showed minor (three or twelve) nucleotide differences (Table 1). *Aspidisca* cf. *cicada* (Figs. S1A, B; 2Q), *B*. *subtropica* (see below), *H*. *teres* (Fig. S1R), *Pseudochilodonopsis* cf. *mutabilis* (Figs. S1C, D; 2P), and *T*. *similis* (see Liao et al., 2021) were identified through both molecular (Table 1) and morphological (on living and stained material) characterizations. Even though we failed to obtain the 18S rRNA gene sequence of *A*. *uncinata* and *P*. *libera*, they could still be identified based on some important morphological characters: for example, *A*. *uncinata* was identified by having the “rolled up” mouth seam overlapping anteriorly, two spherical macronuclei, and few recognizable somatic kineties (Fig. S2C); *P*. *libera* was identified by having one contractile vacuole in the adult and 11–16 cyst ribs (Fig. S2H–K).

*Euplotes* sp. 1 (Fig. S1F–H), *Euplotes* sp. 2 (Fig. S1J–L), *Metopus* sp. (Armophorea, Metopida) (Fig. S1N), *Phialina* sp. (Litostomatea, Haptoria) (Fig. S1O–Q), *Uronema* sp. (Oligohymenophorea, Scuticociliatia) (Fig. S1E, I, M), and *Zosterodasys* sp. (Phyllopharyngea, Synhymenia) (Fig. S2A, B, G) have been identified to genus level based on partial or complete morphological data plus their 18S rRNA gene sequences.

In addition, although we successfully obtained the 18S rRNA gene sequences of two hypotrichous ciliates called Unknown hypotrich sp. 1 and Unknown hypotrich sp. 2, they could not be identified to the genus level because: (1) in general, this molecule is not resolutive for species identification in this group (Spirotrichea, Hypotrichia) due to the numerous polyphyletic genera; (2) a complete morphogenesis data set of Unknown hypotrich sp. 1, which possesses an *Oxytricha*-like 18 frontoventral transverse cirral pattern, is at present lacking (Fig. S2E, F); and (3) unfortunately, only an insufficient number of *in vivo* pictures are available for Unknown hypotrich sp. 2 (Fig. S2D).

### General overview on WWTP ciliate community and its composition

Among the 21 taxa identified above, five species, i.e., *A*. *uncinata*, *P*. *libera*, *Metopus* sp., *Uronema* sp., and *Zosterodasys* sp., were found in original samples only after food enrichment and, in the case of *Metopus*, only after providing strictly anaerobic conditions. The remaining 16 taxa were present in the samples on the days of collection and their relative abundance scores at each sampling time (referred to as □–□□) are shown in Table 1. This matrix has been used to draw the histogram (Fig. 1) where ciliate community’s changes are highlighted especially in relation to functional and trophic categories.

**Figure 1.**
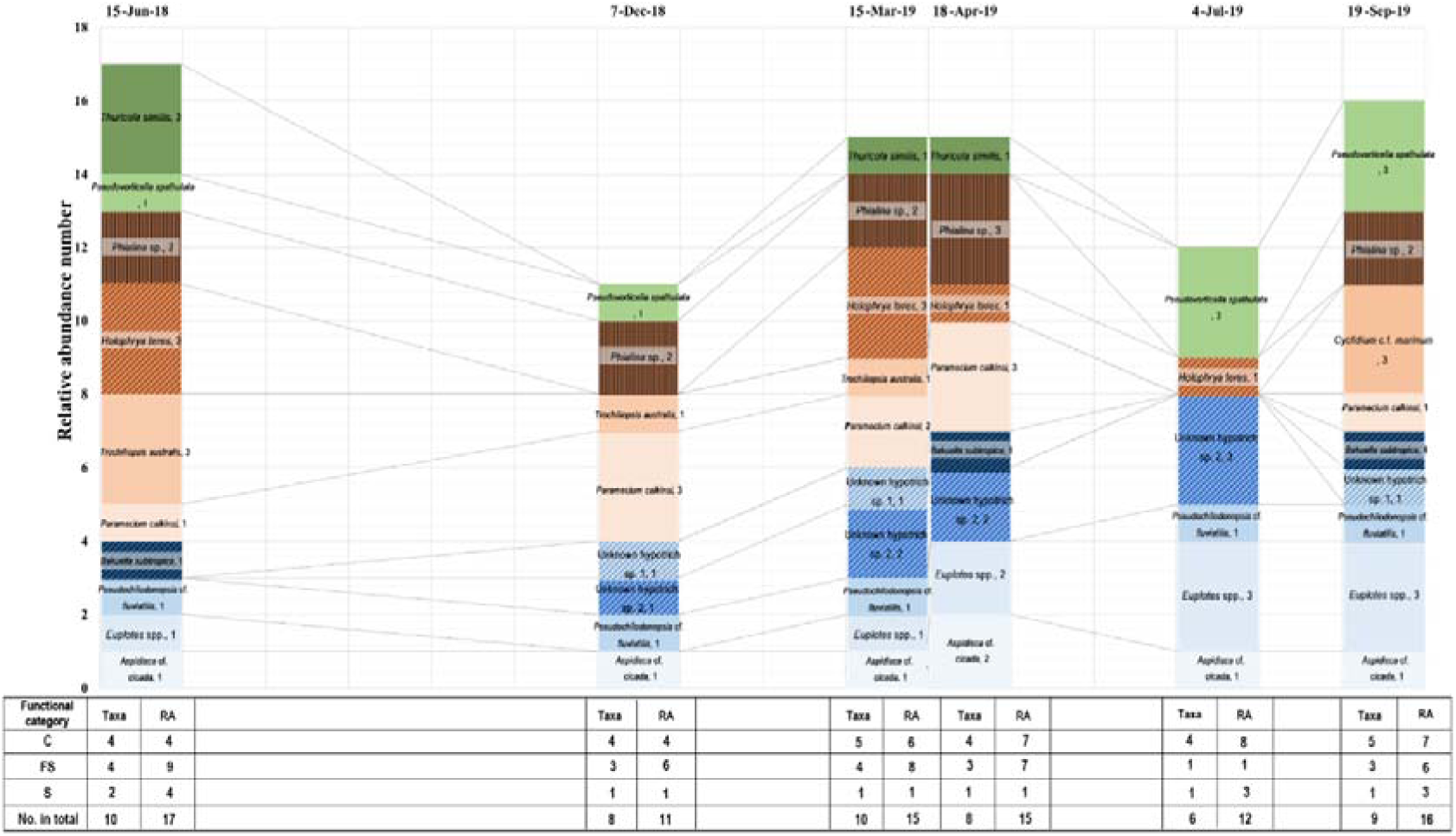
Ciliate community composition at sampling day during the investigation period with the relative abundance and distribution of the ciliates in each sampling occasion (richness table), according to three functional groups. Note that data in this figure are derived from Table 1. In the figure, different background color gradients were used to represent different taxa belonging to the same functional group: blue gradients represent taxa in the “crawling ciliates” group; orange gradients represent taxa in the “free-swimming ciliates” group; green gradients represent taxa in the “sessile ciliates” group. In addition, the trophic category of each species is also indicated in the figure: columns with oblique diagonals represent omnivores; columns with vertical lines represent carnivores; columns without any decoration represent filter-feeding species. Species name plus their relative abundance score (see Table 1 for details) are also indicated on each column. Each white column in the background corresponds to a month. C, crawling ciliates; FS, free-swimming ciliates; No., number; RA, relative abundance score of each functional group; S, sessile ciliates.

Throughout the entire sampling period, the composition of the ciliate community was relatively stable as most of those 16 taxa showed a moderate frequency of occurrence (Table 1). Among them, *Aspidisca* cf. *cicada* was found in all six samplings showing the highest frequency (100%); *Euplotes* spp., *Phialina* sp., *Pseudochilodonopsis* cf. *mutabilis*, and *P*. *calkinsi* were found five times out of six samplings, showing the second highest frequency (83.3%); in contrast, *Cyclidium* cf. *marinum* occurred only once out of six samplings showing the lowest frequency (16.7%) (Table 1). On the other side, only the following species have been regarded to be dominant as for the number of individuals (i.e., ciliates present in massive numbers, whose relative abundance has been marked as “3”) at least in one sampling date: (1) *Euplotes* spp., *H*. *teres*, *P*. *calkinsi*, and *P*. *spathulata* were dominant on two sampling dates each (in detail: *H*. *teres* on I and □□□, *P*. *calkinsi* on II and I□, *Euplotes* spp. and *P*. *spathulata* on □ and □□); (2) *Cyclidium* cf. *marinum*, *Phialina* sp., *T*. *australis*, *T*. *similis*, and Unknown hypotrich sp. 2 were dominant only on a single sampling date, i.e., □□, □□, □, □, and □, respectively (Table 1; Fig. 1).

As for the functional groups, the crawling one was predominant in terms of number of different taxa (Table 1; Fig. 1), even though the four *Euplotes* species were regarded as one systematic/taxonomical unit, as they were so morphologically similar that they could not be properly discriminated based on living observations at the initial stage. Across time, the crawling group included three hypotrichs (i.e., *B*. *subtropica*, Unknown hypotrich sp. 1, and Unknown hypotrich sp. 2), two euplotids (*Aspidisca* cf. *cicada*, and *Euplotes* spp.), and one cyrtophorid ciliate (*Pseudochilodonopsis* cf. *mutabilis*). Free-swimming ciliates were represented by five species: *Cyclidium* cf. *marinum*, *H*. *teres*, *P*. *calkinsi*, *Phialina* sp., and *T*. *australis*. Only two sessile peritrichous taxa were found in this habitat, i.e., *P*. *spathulata* and *T*. *similis*. From the other side, considering the trophic categories, we recorded only one true predator (namely *Phialina* sp.), four “omnivorous” species (i.e., the three crawling hypotrichs plus the free-swimming *H*. *teres*), and 11 remaining, exclusively filter-feeder ciliates. Besides, the carnivorous preference of *Phialina* and the omnivorous behavior of ciliates feeding on other filter-feeders were disclosed by our follow-up observation process based on the residues of some filter-feeders found inside the food vacuoles of these organisms, which also confirmed previous literature records (see Discussion).

### Dynamic analysis of ciliate community based on functional and trophic groups

Although the specific species that constitute the ciliate community were different at the different sampling dates (Table 1, Fig. 1), during the sampling period, the proportions of the three functional groups representing the ciliate community were somehow stable (i.e., four crawling, three free-swimming, and one sessile species). The only exception was the sampling □, where only a single free-swimming ciliate was recorded, and the total number of functional species was also the lowest (Fig. 1). In terms of trophic groups, the secondary consumers (predators or omnivorous) were always present in each sample represented by two to four species (Fig. 1).

The following clues were found during our sampling process when we combined the dynamic changes of each species with its own abundance and functional/trophic classifications (Fig. 1): (1) except for sampling □, the two peritrichs did not appear at the same time, indicating a possible, at least partial, ecological competition between them; (2) *Phialina* sp. was observed on each sampling time with the exception of □ in which the number of species and the relative abundance of the free swimming ciliates was the lowest; (3) for filter-feeding free-swimming ciliates, in general, when *P*. *calkinsi* was present and abundant in number, *T*. *australis* was not and vice versa; when *Cyclidium* cf. *marinum* was present and abundant, *T*. *australis* was not, while *P*. *calkinsi* was present but showing a relatively low abundance.

### Description of Bakuella subtropica based on the Italian population

#### Voucher material

Two voucher slides (registration number-srn: CAMUS_2021-1 and CAMUS_2021-2) with protargol stained cells were produced for *B*. *subtropica* and deposited in the collection of the Museo di Storia Naturale dell’Università di Pisa (Calci, Pisa, Italy) with a black circle on the cover glass showing the position of specimens.

#### General morphology

Cell outline elongate with both ends rounded and posterior slightly narrowed (Fig. 2A–C). Body flexible and slightly contractile, dorsoventrally flattened (Fig. 2D–G). Cell size 72–160 × 28–54 μm *in vivo* (average 135 × 40 μm) (n = 16) and 108–168 × 38–69 μm after protargol staining; length to width ratio between 3:1 and 4:1 *in vivo* (Table 2). Single contractile vacuole located near of left margin, in the middle of the body (Fig. 2C). Pellicle thin and soft, with spherical yellowish cortical granules (about 0.5–1 μm in diameter) distributed on both ventral and dorsal side (Fig. 2F–I). Cortical granules arranged along cirral rows and dorsal bristles, but also distributed as groups or rows in between (Fig. 4H, I). Cytoplasm colorless, packed with some lipids, globules (about 1μm in diameter) and large food vacuoles (about 10–20 μm in dimeter) containing ingested bacteria and small ciliates, like *Aspidisca* cf. *cicada* and/or *Pseudochilodonopsis* cf. *mutabilis*, rendering the cell opaque and dark at low magnification (Fig. 2B–E, P, Q). Many (41–83), ellipsoid to spherical, macronuclear nodules, 2–7 × 1–3 μm in size after protargol staining, scattered throughout the cytoplasm (Table 2; Fig. 2J–M). Micronuclei hard to be recognizable *in vivo*, while two to eight (average: three) micronuclear nodules visible after protargol staining, oval to long elliptical in shape, about 1.7 × 0.9 μm in size (Table 2; Fig. 2M, N).

**Figure 2.**
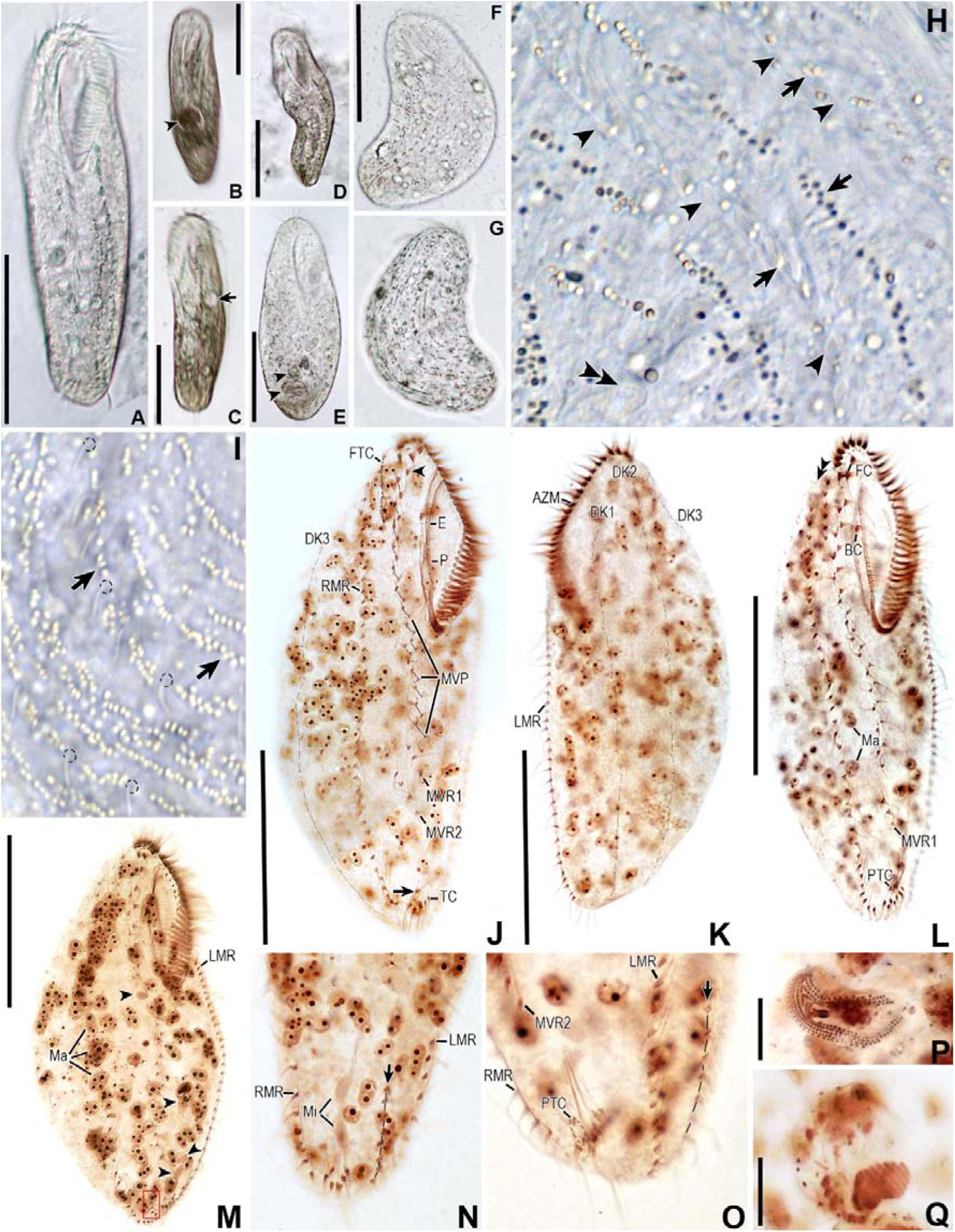
Photomicrographs of *Bakuella subtropica* Italian population *in vivo* (A–I) and after protargol staining (J–Q). A. Ventral view of a representative cell. B–D. Ventral views of other individuals to show a large food vacuole (arrowhead in B) and a single contractile vacuole (arrow in C), as well as the cell flexibility (D). E. Another slightly compressed cell with food debris (arrowheads). F, G. Distribution of cortical granules on the ventral (F) and dorsal (G) side. H. Cortical granules (arrows) arranged near ventral cirral bases (arrowheads) or irregularly distributed in short lines elsewhere. Double arrowhead indicates a macronuclear nodule. I. Cortical granules (arrows) near the dorsal bristles (circular dotted black line) or arranged as short lines in between the rows of bristles. J, K. Composite picture of a protargol stained representative in ventral (J) and dorsal (K) view respectively to show the ciliature on both sides. Arrow indicates a pretransverse cirrus and arrowhead notes the parabuccal cirrus. L. Composite ventral picture of another specimen with only one midventral row, one pretransverse cirrus, and seven transverse cirri. Double arrowhead indicates two pairs of basal bodies located anterior of the right marginal row. M. Ventral view of a specimen with an extra short row (the red rectangle) at posterior end, between transverse cirri and the right marginal row. Arrowheads indicate four elliptical micronuclear nodules. N, O. Ventral views of the posterior portion of two individuals showing an extra short row (arrow) located between transverse cirri and the left marginal row in (N), or on the left side of the left marginal row in (O), respectively. P, Q. Partial ciliature of a digested *Pseudochilodonopsis* cf. *mutabilis* (P) and *Aspidisca* cf. *cicada* (Q) in two different specimens. AZM, adoral zone of membranelles; BC, buccal cirri; DK1–3, dorsal kinety 1–3; E, endoral membrane; FC, frontal cirri; FTC, frontoterminal cirri; LMR, left marginal row; Ma, macronucleus; Mi, micronucleus; MVP, midventral pair; MVR1, 2, midventral row 1, 2; P, paroral membrane; PTC, pretransverse cirri; RMR, right marginal row; TC, transverse cirri. Scale bars: 60 μm (A–F; J–M), 10 μm (P, Q).

**Table 2.**
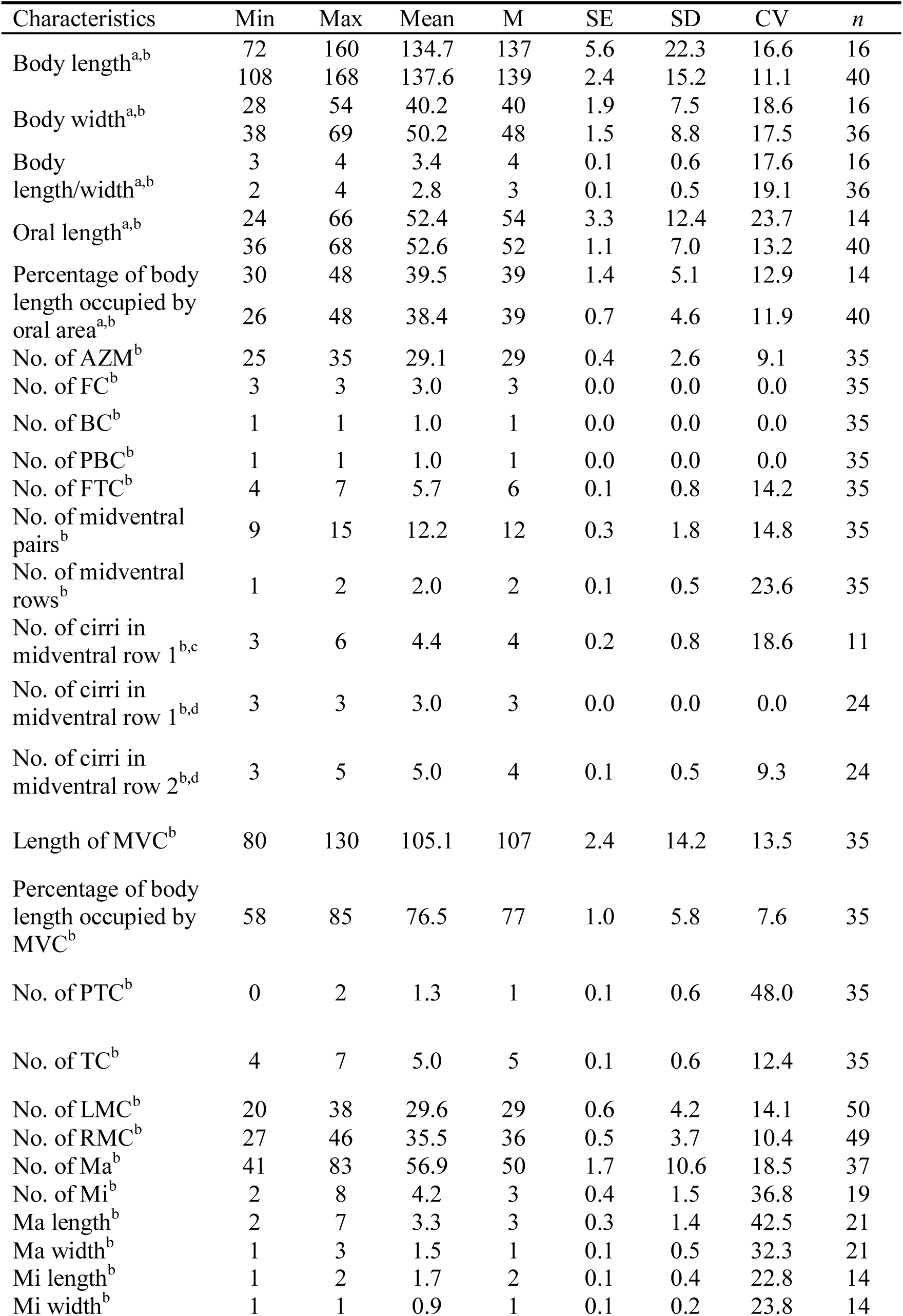
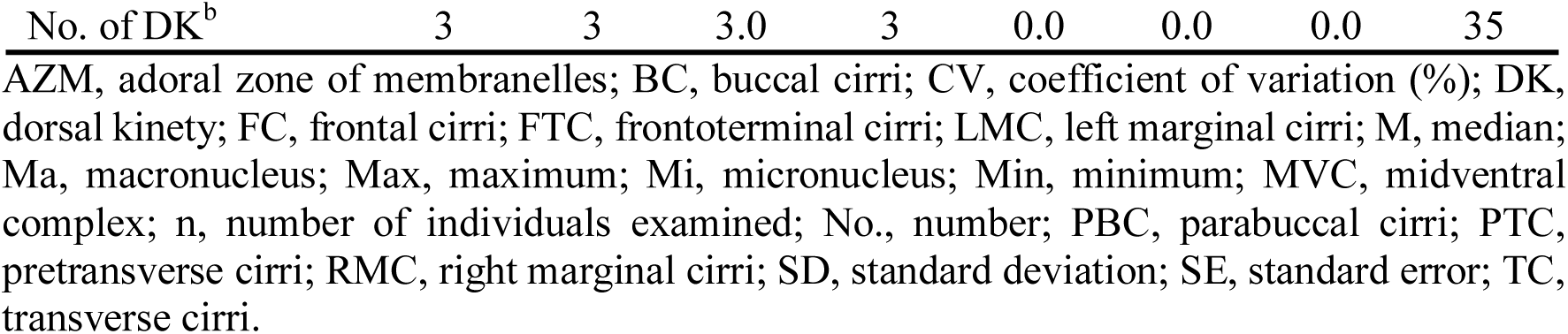
Morphometric data for *Bakuella subtropica* Italian population from living cells (a, first line) and protargol stained specimens (b, second line). c, cells with only one middle ventral cirri row; d, cells with two middle ventral cirri rows. Measurements are in μm.

Adoral zone about 39% of body length, comprising 25–35 membranelles, with cilia 12–17 μm long *in vivo*; buccal cavity deep and moderately wide (Table 2; Fig. 2A–E, J, L, M). Endoral and paroral membranes almost equal in length, distinctly curved, and optically intersected with each other (Fig. 2J, L, M).

Most somatic cirri relatively fine with cilia about 10 μm long *in vivo* (Fig. 2A). Consistently three, relatively stout, frontal cirri, one parabuccal cirrus, and one buccal cirrus near paroral membranes (Table 2; Fig. 2J, L, M). Four to seven frontoterminal cirri near distal end of adoral zone (Table 2; Fig. 2J, L, M). Midventral complex composed of 9–15 pairs of cirri arranged in the typical Urostylida-like zig-zag pattern, following by one or two short midventral rows: 11 out of 35 observed specimens only possessing a single midventral row composed of three to six cirri (Table 2; Fig. 2L, M); the remaining 24 showing two midventral rows (with the anterior one consistently comprising three cirri) and the posterior one composed of three to five cirri (Table 2; Fig. 2J). Such structure generally extending to about 77% of body length (Table 2). Four to seven (usually five) transverse cirri located near posterior end of cell with cilia about 15 μm long *in vivo* and projecting beyond the rear body end (Table 2; Fig. 2J, L, M–O). A single (in 18 out of 35 observed specimens) or two (in 14 out of 35 observed specimens) pretransverse cirrus/i also observed in most protargol stained cells with relatively fine bases respect to those of transverse cirri (Table 2; Fig. 2J, L, M–O). One left and one right marginal row comprising 20–38 and 27–46 cirri, respectively (Table 2; Fig. 2J–M). Three bipolar dorsal kineties with bristles about 3 μm long *in vivo* (Table 2; Fig. 2I, K). Two additional dikinetids usually present at anterior part of right marginal cirral row (Fig. 2L). Caudal cirri absent (Fig. 2K). Noteworthy, in a few individuals, an extra cirral row at posterior end, specifically located between the transverse cirri and the marginal rows (Fig. 2M, N), or an additional short marginal row located on the left rear side of the left marginal cirral row were observed (Fig. 2O). However, those few individuals still possessed the same ciliature pattern as described before, so the abovementioned extra cirral row has not been included in Table 2. Locomotion by moderately crawling on debris, sometimes swimming, and slowly rotating around the main body axis.

#### Ribosomal gene sequence and analysis

The full ribosomal operon of *B*. *subtropica* Italian population is 5,165 bp long excluding PCR primers and includes an overall GC content of 44.86%. In GenBank its accession number is MZ067023 and it comprises the almost complete 18S rRNA (1,750 bp), the ITS1 (132 bp), the 5.8S (152 bp), the ITS2 (191bp), and the almost complete 28S rRNA (2,940 bp) sequences. In terms of 18S rRNA gene, our organism is 100% identical to the Korean population of *Bakuella incheonensis* (KR024011, 1746 bp) and the Chinese population 2 (pop 2) of *B*. *subtropica* (KY874001, Shanxi population, 1721 bp), but the sequence of the type Chinese population of *B*. *subtropica* (pop 1, Guangzhou population), KC631826, deviates from all mentioned three sequences in the same two positions. Moreover, concerning the ITS and 28S rRNA gene, the Italian population shows only a one-nucleotide difference from KY874003 (ITS gene sequence of *B*. *subtropica* pop 2) and a five-nucleotide difference from KY874002 (28S rRNA gene sequence of *B*. *subtropica* pop 2). Unfortunately, the ITS and 28S rRNA gene sequences of *B*. *incheonensis* are unavailable at present.

#### Mitochondrial genome

The linear mitochondrial assembly of *B*. *subtropica* was 52,184 bp in length, with a GC content of 31.9%. It was deposited in the GenBank database with the accession number MZ292454. Its gene content was composed by 39 open reading frames (ORFs), a 12S rRNA gene (*rns*), a partial 16S rRNA gene (*rnl*_*a*), and 13 tRNA genes (Fig. 3). 23 out of 39 retrieved ORFs were protein coding genes, namely *nadh1*_*a*, *nadh2*_*a*, *nadh2*_*b*, *nadh4*, *nadh5*, *nadh6*, *nadh7*, *nadh9*, *nadh10*, *rpL2*, *rpL6*, *rpL14*, *rpS3*_*b*, *rpS4*, *rpS7*, *rpS10*, *rpS12*, *rpS13*, *cob*, *cox1*, *cox2*, *ccmf*_□, and *ccmf*_□. The remaining 16 ORFs are unclassified with unknown function. Besides, there was a AT repeat region in the middle of the mitogenome of *B*. *subtropica*, made up of 13 tandem repetition of TAATTAATT[TA]_n_CGTATAT with a variable number of TA dimers spanning from four to seven, working as a bi-directional transcription start.

**Figure 3.**
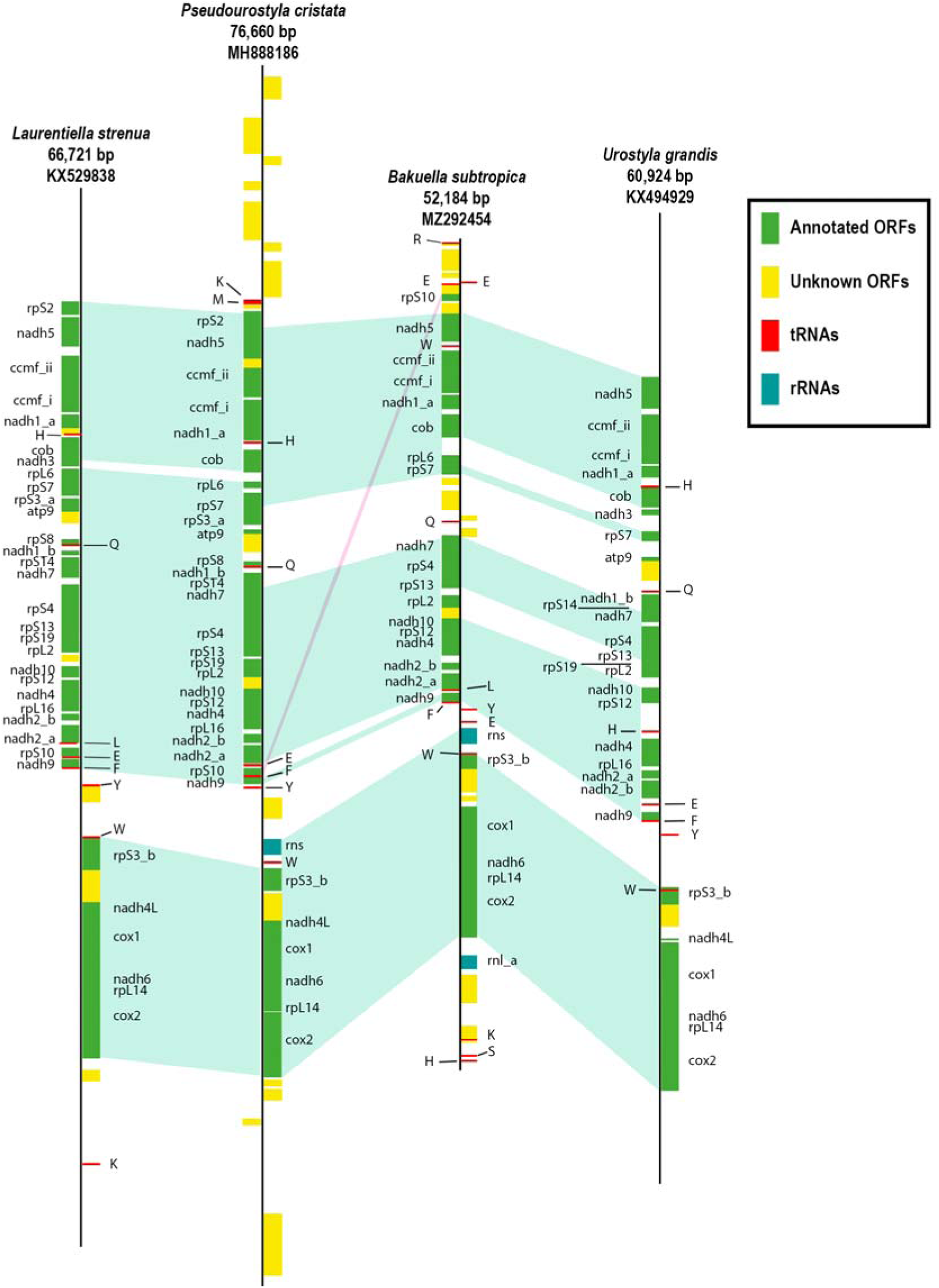
Mitochondrial gene map of *Bakuella subtropica* and some representatives within the subclass Hypotrichia, with the GenBank accession number attached. Split genes are suffixed with an underscore followed by an alphabetic character; the putative split gene, namely *ccmf*, is suffixed with an underscore followed by a lowercase roman numeral. tRNAs are indicated by a single letter according to amino acid codon table. Green pale areas indicate syntenic regions between the analyzed genomes, while the structural rearrangement is indicated by pale-pink shade.

#### Phylogenetic and phylogenomic analyses

Phylogenetic trees inferred from the 18S rRNA gene sequences, using two different methods (namely, maximum likelihood (ML) and Bayesian inference (BI)) showed similar topologies, therefore, only the ML tree (Fig. 4) was presented with bootstraps and posterior probabilities from both algorithms. In the phylogenetic tree, *B*. *subtropica* Italian population (MZ067023) was placed within the core Urostylida and clustered together with other two *B*. *subtropica* sequences (KY874001 and KC631826) as well as the one referred to as *B*. *incheonensis* (KR024011), forming a polytomy with strong support (96/0.99). Then it clustered with *Anteholosticha paramanca* (KF806443) (90/1.00), forming a sister group to the clade composed by other two *Anteholosticha* spp., namely *Anteholosticha manca* (DQ503578) and *Anteholosticha multicirrata* (KC307773) (56/0.72).

**Figure 4.**
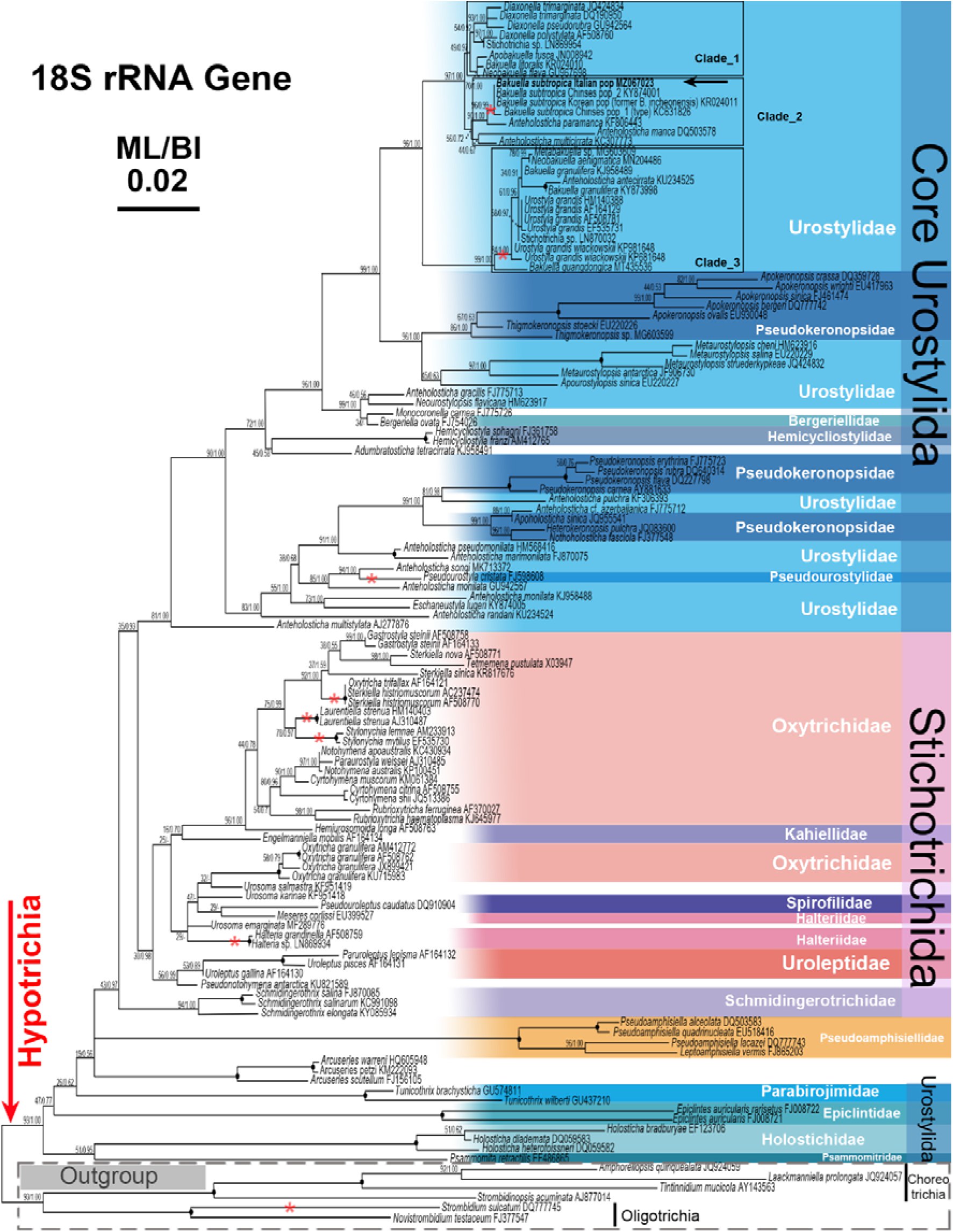
Phylogenetic tree inferred from 18S rRNA gene sequences showing the position of *Bakuella subtropica* Italian population (bold font, arrowed). Support values near nodes are bootstrap values for ML and posterior probabilities for BI, respectively. Black dots indicate nodes with full support in both analyses. Clades with a different topology in the BI tree are indicated by “-”. Red asterisks indicate the closest branches that contain the species used also in subsequent phylogenetic analysis (Fig. 5) based on mitochondrial genome. All branches are drawn to scale. GenBank accession numbers are given for each species. The scale bar represents two substitutions per 100 nucleotide positions. Clades 1, 2, and 3 represent the clades of *Bakuella*-related species.

The ML phylogenetic tree based on mitochondrial genes (Fig. 5) showed a topology almost concordant with that inferred from 18S rRNA gene data (Fig. 4) in terms of the representatives of the subclasses Oligotrichia and Hypotrichia, with the exception of *Pseudourostyla cristata* that did not cluster together with the other two urostylids (*B. subtropica* and *Urostyla grandis*) but branches as a sister taxon to Stichotrichida.

**Figure 5.**
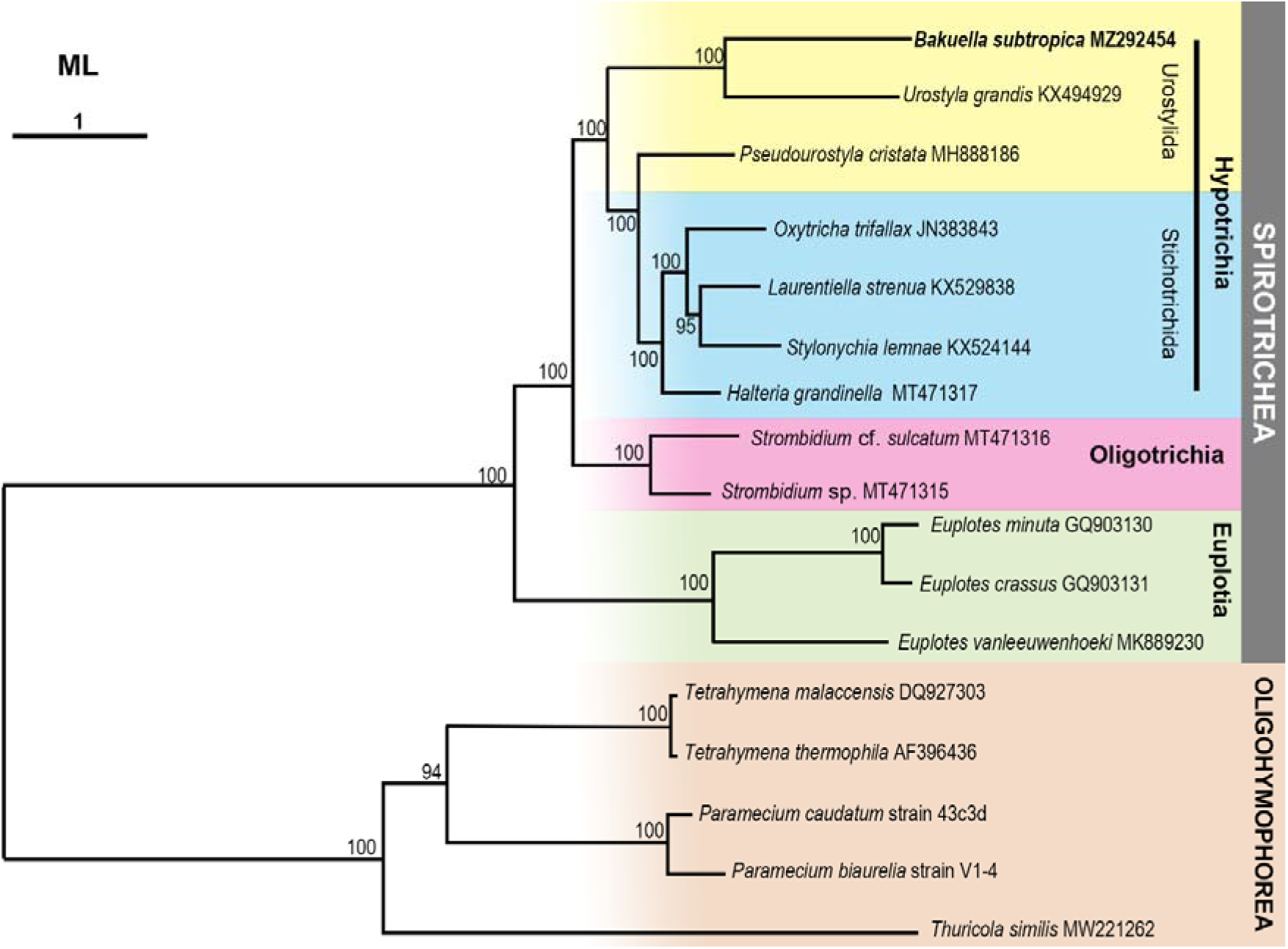
ML phylogenetic tree of *Bakuella subtropica* (bold font) inferred from the concatenated amino acid sequences alignment of 18 selected mitochondrial protein coding genes. Numbers on each node represent bootstrap values. The scale bar (1) denotes mean number of substitutions per site. Entries without accession numbers were downloaded from ParameciumDB database.

## Discussion

### Identity of the ciliate species found in Cuoiodepur WWTP

Since 1970s, study on ciliates from WWTPs has bloomed, hundreds of papers on faunistic, ecological, and taxonomy have been published (e.g., Curds and Cockburn, 1970; Curds, 1973a, b; Esteban et al., 1991; Aescht and Foissner, 1992; Madoni et al., 1993; Salvadó and Gracia, 1993; Madoni, 1994; Salvadó, 1994; Martín-Cereceda et al., 1996; Lee et al., 2004; Liu et al., 2008; Pérez-Uz et al., 2010; Dubber and Gray, 2011; Madoni, 2011; dos Santos et al., 2014; Matsunaga et al., 2014; da Silva Paiva et al., 2016; Foissner, 2016; Chouari et al., 2017), giving us a detailed understanding on some organisms frequently found in this environment and their possible roles in the wastewater treatment process. However, in terms of species identification, most of faunistic and ecological works were only based on *in vivo* sample check under the microscope with the aid of some specific manuals (e.g., Curds and Cockburn, 1970; Esteban et al., 1991; Madoni et al., 1993; Martín-Cereceda et al., 1996; Dubber and Gray, 2011; dos Santos et al., 2014), while few studies used either only the classical taxonomic approach (i.e., a combination of living observation and silver staining techniques) (Aescht and Foissner, 1992; Salvadó and Gracia, 1993; Salvadó, 1994; Pérez-Uz et al., 2010) or only molecular techniques (Matsunaga et al., 2014; Chouari et al., 2017). At the same time, some taxonomical studies (using classical and/or molecular methods) on species found in sewage have also been carried out, showing the uniqueness of this habitat as a source of both poorly known and novel ciliate species (e.g., Foissner et al., 1988; Leitner and Foissner, 1997; Guggiari and Peck, 2008; da Silva Paiva et al., 2016; Foissner, 2016).

In this context, the present work firstly applied the integrative taxonomy (living observation, silver staining techniques, 18S rRNA gene sequencing etc.) to identify all ciliate species found in the activated sludge of a WWTP. Even though, the process is time-consuming and requires different expertise, it definitely provides a more reliable species list and has additional taxonomic values. Moreover, we also performed further detailed descriptions of some retrieved species according to NGTax, combining mitogenome-based phylogenomic analysis with integrative taxonomy approach. From this perspective, we recently redescribed and neotypified the poorly known peritrich *T. similis*, collected from Cuoiodepur WWTP (Liao et al., 2021), while in the present work, we provide the characterization of *B. subtropica*, a ciliate presenting a conflict in species identification. Our improved description is based on a multidisciplinary study, where results obtained from the different applied analyses have been compared and integrated (see below). Finally, two among the characterized species, i.e., *P*. *calkinsi* and *P*. *spathulata*, isolated from Cuoiodepur WWTP, had been previously used in batch experiments aimed at evaluating the role of bacterivorous organisms on fungal-based systems for natural tannin degradation (Sigona et al., 2020). Intriguingly, *P*. *spathulata*’s 18S rDNA sequence characterized by Sigona et al. (2020) (MT025819) differs from the currently studied population (MZ098633) by possessing a rather long (>900 bp) intron, while the remaining part of the two sequences are coincide.

### General overview on ciliate community and its composition

One of the most used techniques in wastewater treatment is biological treatment with activated sludge, in which a complex microbiological community is suspended and plays the key role for organic compounds and nutrients removal (Comeau, 2008). It has already been observed that the taxonomic composition and population distribution of protozoa in the activated sludge system are directly related to (1) the nature (i.e., domestic, textile, leather, or dairy) of the wastewater involved by concentrations of organic and inorganic nutrients; (2) the operating conditions, such as aeration intensity, solids retention time (SRT), hydraulic retention time (HRT), temperature, and pH; (3) and the system configuration, specifically, different combinations of one or several types of reactors (i.e., aerobic, anoxic, and anaerobic), and the recirculation of activated sludge between them on the system pipeline to meet different requirements (e.g., Esteban et al., 1991; Salvadó and Gracia, 1993; Salvadó, 1994; Martín-Cereceda et al., 1996; Lee et al., 2004; Liu et al., 2008; Dubber and Gray, 2011; dos Santos et al., 2014).

In our specific case, we mostly dealt with industrial wastewater: in fact, about 50% of flow rate (and 97% of organic load) derived from tannery industry and 50% of flow rate (and 3% of organic load) from domestic wastewater. Cuoiodepur applies MLE process, which consists of equalization, primary settling, predenitrification (anoxic) and nitrification (aerobic) biological section (where most of nitrogen and organic load is removed), and a tertiary treatment to achieve high quality effluent standard before the discharge in the receiving water body (the Arno River). Moreover, it is worth remembering that Cuoiodepur wastewater has salinity in the range of 5‰ to 10‰, indicating that it is a brackish environment. In this context, the occurrence of in total 21 taxa is somehow consistent with previous studies on ciliated protozoa community in other activated sludge plants applied oxic/anoxic configures (Liu et al., 2008; Pérez-Uz et al., 2010; Dubber and Gray, 2011).

Most taxa retrieved in Cuoiodepur have been frequently recorded in other WWTPs in different geographic and climatic locations (as examples, see Curds and Cockburn, 1970; Foissner and Berger, 1996; Guggiari and Peck, 2008; Madoni, 2011; Foissner, 2016)), but the studied WWTP also showed peculiarities, with the presence of some rarely recorded taxa, like *B*. *subtropica*, *Phialina* sp., *T*. *australis*, *T*. *similis*, and *Zosterodasys* sp. Among them, *T*. *australis* and *T*. *similis* have their original type-locality in sewage (Bock, 1963; Foissner et al., 1988). *Phialina* and *Zosterodasys* are sometimes reported as rather abundant in high-load and/or oxygen deficient activated sludge (Foissner and Berger, 1996; Foissner, 2016). As sewage is a recently established artificial habitat, we can expect that the species found have their own natural habitat. In this regard, it is intriguing to observe that *B*. *subtropica* has originally been described in estuarine and nutrient rich brackish environment (Chen et al., 2013), while another population of *T*. *australis* was found by Fan et al. (2014) in a brackish mangrove wetland.

Moreover, apart from the species retrieved in the samples at collection days, we also recorded some species appearing later during laboratory maintenance. This situation was probably due to ciliates’ encystment and excystment ability, with the excystment occurring in more favourable conditions, such as nutrient availability and aerobic/anaerobic requirements. Although those later-appearing species might not be able to participate in the composition of the real community in the WWTP, and our data did not support a relevant ecological role for these organisms, there are several worth noticing points: (1) the development of *A. uncinata* is determined by the quality of the sewage influent (Martín-Cereceda et al., 1995), thus the industrial wastewater with high chemical oxygen demand consumption (Table S2) in Cuoiodepur may not favour the presence of high density of this organism; also, its small, flat, and transparent body (Table 1; Fig. S2C) and its behaviour of attaching on sludge flocs, should be considered, as they make the cell particularly difficult to observe in samples. (2) *Uronema* is a small, fast growing organism, representative of the free-swimming bacterivorous ciliates linked to the first phase of colonization of the plant; it usually does not appear or become dominant in a WWTP at the steady state, and its sudden demographic bloom is usually considered as an indicator of deteriorated condition resulting from disfunctions (Madoni, 2011; Foissner, 2016). (3) *P*. *libera* was recorded once but at low numbers by Holm (1925) in Elbe estuary near Hamburg, where endured huge volumes of wastewater. Compared to free-swimming carnivores, *Podophrya* is rarely recorded, possibly due to its small size and encystment capability. (4) A “true” anaerobic ciliate, *Metopus* sp. was exclusively present in the tightly sealed sampling bottles collected from the denitrification tank after one or two months of cultivation, but it had never been found in freshly collected samples. In this regard, also considering that no significant differences in the ciliate community structure were found between the aerobic and the anoxic tanks at the sampling day, there are indications that (a) species retrieved at the sampling day are able to survive the oxic/anoxic cycles typical of the studied system and are probably selected by such cycling conditions; (b) the conditions for the mass growth of obligate anaerobic ciliates are very strict, and the high recirculation between aerobic and anoxic tanks in this system does not allow strict anaerobic species to proliferate.

During our sampling, 15 out of 16 species were retrieved with a moderate frequency (≥ 50%) (Table 1), indicating that the operation of Cuoiodepur WWTP along the year was basically stable and the slight fluctuations of physico-chemical parameters of the week before sampling (Table S3) might have an impact on the species composition collected the following week, without affecting the overall species composition of the WWTP. The species with high frequency (≥ 83.3%) and relative abundance (score ≥ 2), namely *Aspidisca* cf. *cicada*, *Euplotes* spp., *P*. *calkinsi*, and *Phialina* sp. (Table 1), should be consequently considered as typical representatives of the ciliate community of Cuoiodepur. Among them, *Aspidisca* cf. *cicada*, *Euplotes* spp., and *P*. *calkinsi* are known to have strong ecological tolerance, e.g., they can survive in highly polluted water bodies (Madoni et al., 1992; Madoni et al., 1994; Puigagut et al., 2005; Sobczyk et al., 2020) and develop high-density populations. In addition, *Aspidisca* cf. *cicada* is one of the most frequently reported species in WWTP receiving industrial inputs (Dubber and Gray, 2011; Sobczyk et al., 2020); its occurrence is in line with the characteristics of the studied plant where the high concentration of ammonia and toxic tannery wastewater is treated. *Phialina* sp., the free-swimming carnivorous ciliate, appeared five times out of six samples; its high occurrence might reflect another characteristic of the studied plant, that is the presence of a well-structured food web (see below).

*Cyclidium* cf. *marinum* only occurred once (i.e., in September 2019) out of six samplings, but was rather abundant (score = 3). Interestingly, August is the traditional period of summer vacation in Italy, and, during this month, most tanneries, whose wastewater is collected in the plant, close, therefore the industrial flow rate is drastically reduced. It has already been proved in Giordano et al. (2016) that the microbial community structure in Cuoiodepur reshaped at this breakpoint, but, unfortunately, that work did not conduct a continuous and stable frequency monitoring on ciliate community. Based on the data from our six samplings, we hypothesise that the sudden outbreak of *Cyclidium* cf. *marinum* observed in the sample of September may be due to the reduced industrial flow at the holiday, which caused the entire working environment of WWTP to become more suitable for the growth of this organism.

### Dynamic analysis of ciliate community based on functional and trophic groups

Competition, predation, and other trophic relationships among the ciliates of the WWTP community, along with plant management practices, lead to a succession of ciliate populations until dynamic stability is reached (Nicolau et al., 2005).

A “well-functioning plant” is characterized by a ciliate community dominated by crawling and sessile species (Martín-Cereceda et al., 1996; Nicolau et al., 2005; Madoni, 2011). We could observe such a structure only on two sampling dates (i.e., V and VI; Fig. 1); additionally, although sometimes the relative abundance of sessile group was rather high, its taxon diversity was unexpectedly low (i.e., only two species), which is also the main reason why crawling and sessile groups do not dominate the community in most of the sampling cases. Also, in Martín-Cereceda et al. (1996), two out of the ten studied WWTPs had a significant abundance of swimming ciliates which may be due to the nature of the received industrial inputs. A dominance of free-swimming ciliates, such as that observed in samples □–□□, is typically considered to be linked to the initial set-up phase of the plant or to short-term disorder phases during the steady period. On the other side, monitoring data of Cuoiodepur’s effluent (Table S2) guarantee that the plant is well performing despite this unusual, in terms of functional categories, ciliates species composition. We suggest two different explanations for this observation: (1) the unique composition of the ciliate community in Cuoiodepur may be attributed to the wastewater nature (tannery), and/or to the configurations used, and/or to other specific plant’s aspects; (2) the plant suffers from periodical spikes of pollutants that perturbate the ciliate community preventing in most of the cases its proper structuring (i.e., with a high number of crawling and sessile ciliates). Only further study will allow to solve the issue.

Moreover, during our sampling process, the seldom co-occurrence observed for the two peritrichous ciliates suggests a kind of competition between them, i.e., they might share partially overlapped niches. The competition is also apparently occurring among some free-swimming ciliates such as the couples represented by *Paramecium*/*Trochiliopsis* or *Trochiliopsis*/*Cyclidium*. In this case, we might speculate that the fast-growing r-selected species, *Trochiliopsis* and *Cyclidium*, can be possibly slowly replaced by *Paramecium* (the k-selected species) when condition gets stabilized.

Another evident feature of the studied ciliated community is the presence of secondary consumers which feed on filter-feeders; they may play a role in modifying bacterivorous species diversity and indirectly influence bacterial communities (Curds, 1973b; Madoni, 1994; Pajdak-Stós et al., 2017). In the system we observed, the free-swimming *Phialina*, a well-known carnivorous ciliate, is a typical representative of secondary consumers, and its presence was always found associated to free-swimming filter-feeders (potential preys), suggesting a direct trophic link that has partially been confirmed by laboratory observation where *Uronema* sp. (a species generally absent in freshly collected samples and appearing in lab after enrichment) was found in food vacuole of *Phialina* (W.L. personal observations). Apart from *Phialina*, other carnivorous representatives of the system are the sessile suctorian *P*. *libera* and the crawling *A*. *uncinata*. Although they were not found on the day of sampling, in some circumstances, these organisms may share the same role as *Phialina* after the excystment. Indeed, in our lab observation, *P*. *libera* was found to feed on some free-swimming ciliates (like *P*. *calkinsi* and small scuticociliates). *A*. *uncinata* is reported to feed on small hymenostome ciliates, e.g., *Cyclidium*, *Uronema*, and flagellates (Augustin et al., 1987). Furthermore, we observed another group of secondary consumers represented by omnivorous ciliates, namely three crawling hypotrichs and the free-swimming *H*. *teres*, all commonly found at the sampling day. They proved to selectively feed on filter-feeders (in details, smaller crawling ciliates, i.e., *Aspidisca* cf. *cicada* and *Pseudochilodonopsis* cf. *mutabilis*, were observed in the food vacuoles of hypotrichs, whereas the free-swimming *P*. *calkinsi*, *Uronema* sp., and the possible swarmer of *P*. *spathulata* were recognizable in food residues of *H*. *teres*), thus probably contributing to regulate filter-feeder populations.

To our best knowledge, despite their practical significances, the interdependencies among activated sludge eukaryotic microorganisms have not been thoroughly investigated up to now. Although still preliminary and requesting further experimental confirmation, the observed trophic relationships, schematized in Fig. 6, start shedding light on trophic web structure in this environment. In this context, it is worth mentioning that, in addition to ciliates, other protists (such as flagellates and amoebae like *Arcella* spp.) and few metazoan taxa (such as rotifers and nematodes) were also observed during the survey: their roles in the food-web of Cuoiodepur WWTP will be addressed in future investigation.

**Figure 6.**
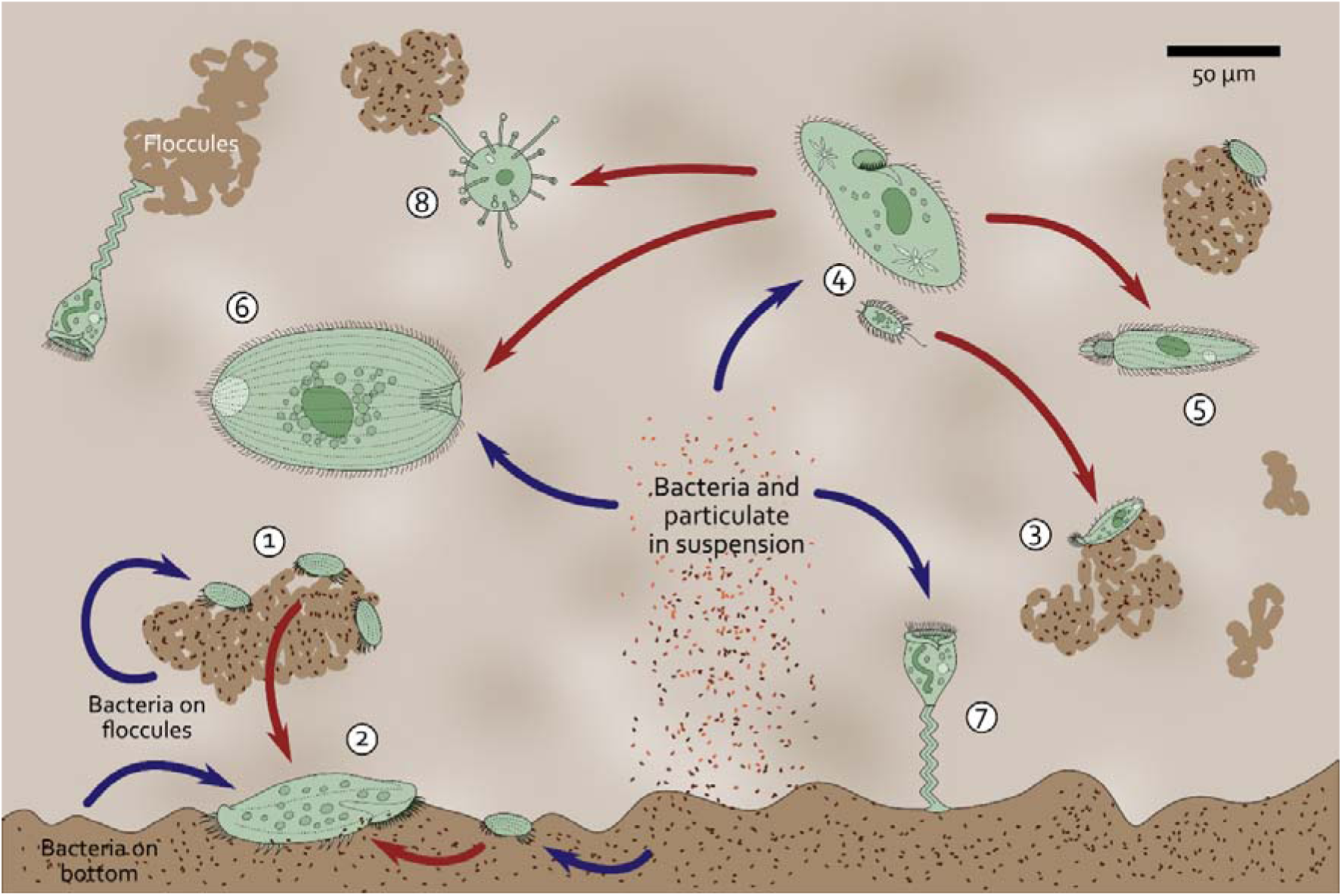
Illustration showing the interrelationships among representatives of the organisms found in the ciliate community of the Cuoiodepur WWTP. 1, Crawling filter-feeders (represented by *Aspidisca*); 2, Crawling omnivores (represented by hypotrichs); 3, Crawling carnivores (represented by *Acineria*); 4, Free-swimming filter-feeders (represented by large *Paramecium* and small scuticociliates, like *Uronema*); 5, Free-swimming carnivores (represented by *Phialina*); 6, Free-swimming omnivores (represented by *Holophrya*); 7, Sessile filter-feeders (represented by *Vorticella*-like species); 8, Sessile carnivores (represented by *Podophrya*). Blue and red arrows indicate bacterivorous and carnivorous feeding behaviours, respectively.

### Bakuella subtropica: proposal of synonymization with Bakuella incheonensis

*B*. *subtropica* (pop 1) was firstly discovered by Chen et al. (2013) from brackish water sample in the estuary of the Pearl River in Guangzhou (China) with detailed morphological, morphogenesis, and phylogenetic descriptions. Later, Jo et al. (2015) discovered a *Bakuella* species in brackish water near Aamdo Shore Park, Incheon, (South Korea), which has been morphologically and molecularly well described. Even though the Korean *Bakuella* sp. and *B*. *subtropica* were extremely similar at a molecular level (showing only a two-nucleotide difference in the whole 18S rRNA gene sequence), they still could be separated based on some morphological characteristics (such as body size; number of adoral zone membranelles, frontoterminal cirri, midventral pairs, left and right marginal cirri, and macronuclear nodules; length of midventral complex; and kind of cortical granules). Hence, Jo et al. (2015) regarded it as a new species, named *B*. *incheonensis*. Subsequently, Lyu et al. (2018) recorded a different *B. subtropica* (pop 2), from Shanxi (China), whose 18S rRNA gene sequence was identical to that of *B*. *incheonensis*, but, unfortunately, a morphological comparison between the two organisms was not possible because no morphological data were concurrently provided. Finally comes our present study on the population of *Bakuella* from Cuoiodepur WWTP: we found that the morphological differences previously used for the discrimination between *B. incheonensis* and *B*. *subtropica* appear inconspicuous when taking our Italian population into consideration, because most of the morphological data overlap in their respective range among the three populations in analysis (Table 3). Moreover, as mentioned before, the 18S rRNA gene sequences of MZ067023 (Italian population), KR024011 (*B*. *incheonensis*), and KY874001 (*B*. *subtropica* pop 2) are 100% identical, and the two-nucleotide difference from the sequence KC631826 (*B*. *subtropica* pop 1) might possibly be due to PCR errors fixed by the cloning approach used by the authors to characterize the sequence. Indeed, these only two sites are located in character columns that are conserved in all organisms of the clade and, consequently, should not be variable at a species level. In summary, aiming to contribute to clarifying the *Bakuella* intrageneric relationships, according to the whole body of available data we propose that *B*. *incheonensis* should be considered as a junior synonym of *B*. *subtropica*. However, it cannot be excluded that *B*. *incheonensis/B*. *subtropica* represents a group of sibling species; unfortunately, molecular data for fast-evolving marker, i.e., mitochondrial genes, of the other *Bakuella* populations are unavailable so far, thus this hypothesis cannot be tested at present.

**Table 3.**
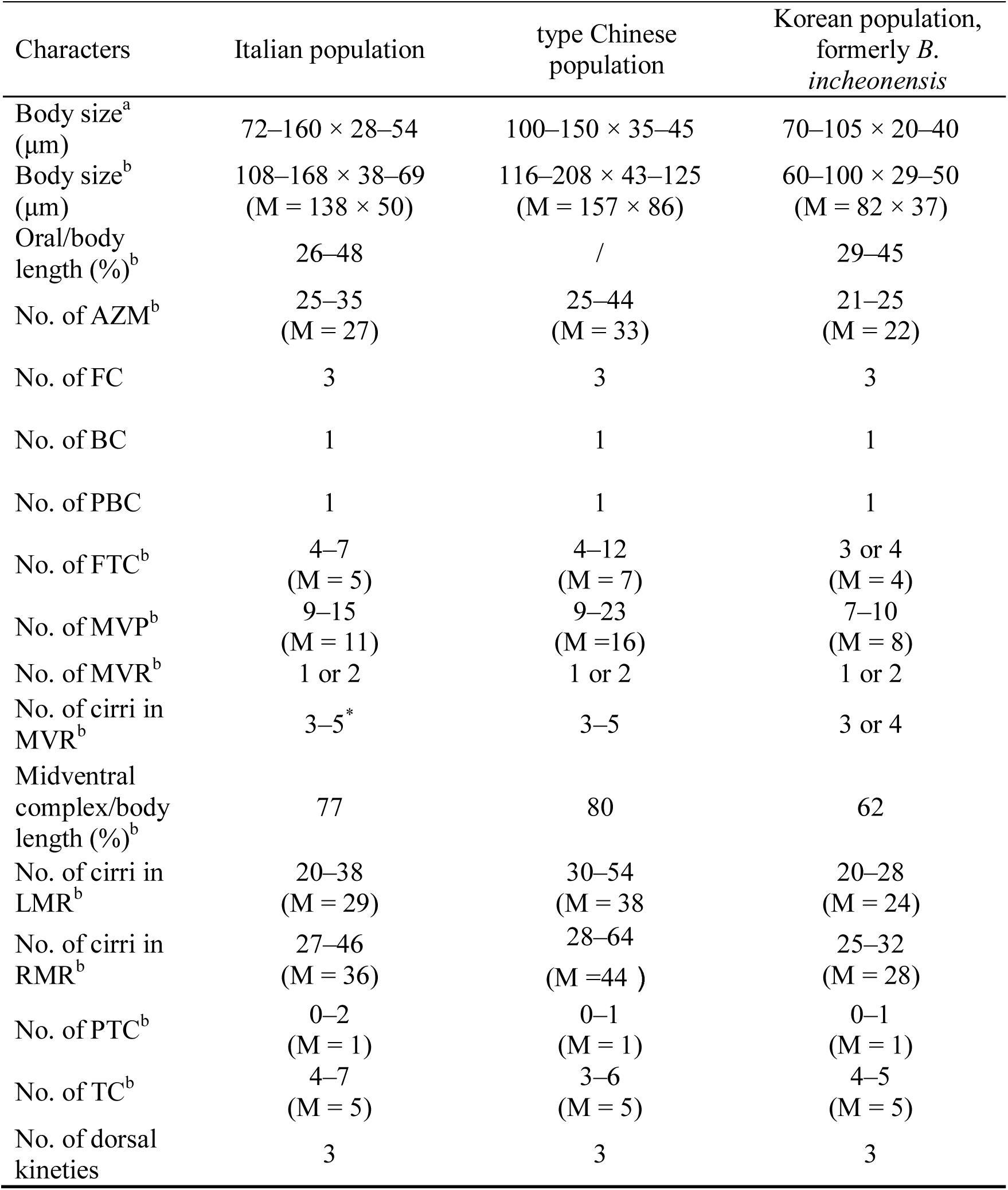
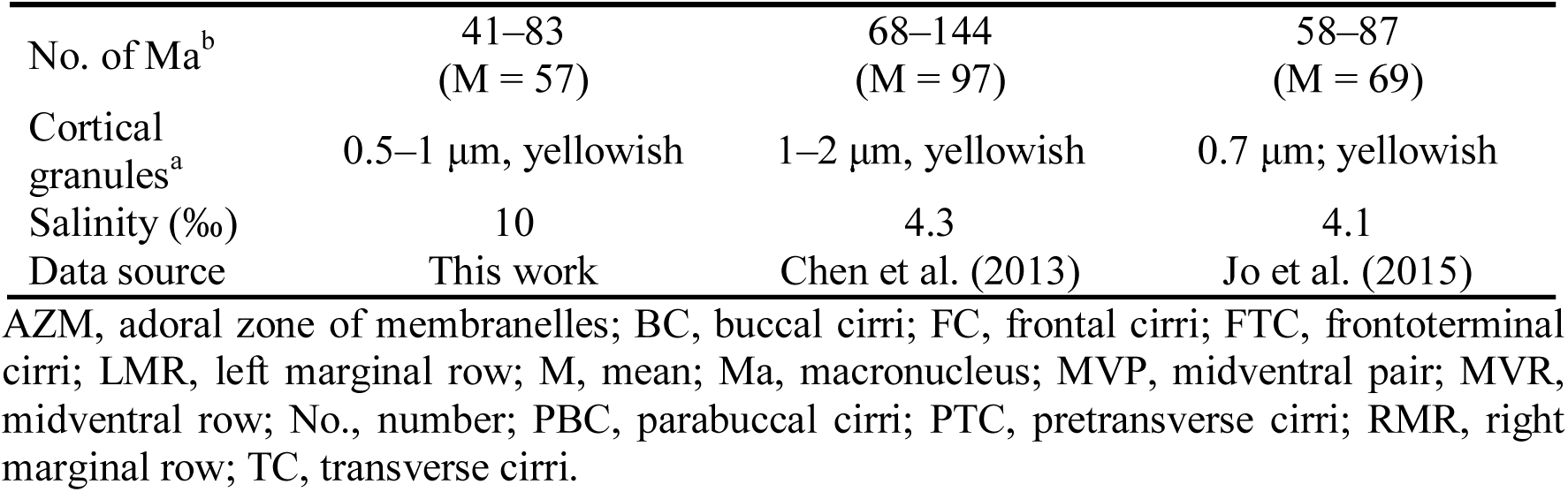
Morphological comparison among *Bakuella subtropica* populations. /, data unavailable; a, data from living cells; b, data from protargol stained specimens; *, data from cells with two midventral rows.

| Characters | Italian population | type Chinese population | Korean population, formerly *B. incheonensis* |
| --- | --- | --- | --- |
| Body size<sup>a</sup><br>(μm) | 72–160 × 28–54 | 100–150 × 35–45 | 70–105 × 20–40 |
| Body size<sup>b</sup><br>(μm) | 108–168 × 38–69<br>(M = 138 × 50) | 116–208 × 43–125<br>(M = 157 × 86) | 60–100 × 29–50<br>(M = 82 × 37) |
| Oral/body<br>length (%)<sup>b</sup> | 26–48 | / | 29–45 |
| No. of AZM<sup>b</sup> | 25–35<br>(M = 27) | 25–44<br>(M = 33) | 21–25<br>(M = 22) |
| No. of FC | 3 | 3 | 3 |
| No. of BC | 1 | 1 | 1 |
| No. of PBC | 1 | 1 | 1 |
| No. of FTC<sup>b</sup> | 4–7<br>(M = 5) | 4–12<br>(M = 7) | 3 or 4<br>(M = 4) |
| No. of MVP<sup>b</sup> | 9–15<br>(M = 11) | 9–23<br>(M = 16) | 7–10<br>(M = 8) |
| No. of MVR<sup>b</sup> | 1 or 2 | 1 or 2 | 1 or 2 |
| No. of cirri in<br>MVR<sup>b</sup> | 3–5* | 3–5 | 3 or 4 |
| Midventral<br>complex/body<br>length (%)<sup>b</sup> | 77 | 80 | 62 |
| No. of cirri in<br>LMR<sup>b</sup> | 20–38<br>(M = 29) | 30–54<br>(M = 38) | 20–28<br>(M = 24) |
| No. of cirri in<br>RMR<sup>b</sup> | 27–46<br>(M = 36) | 28–64<br>(M = 44) | 25–32<br>(M = 28) |
| No. of PTC<sup>b</sup> | 0–2<br>(M = 1) | 0–1<br>(M = 1) | 0–1<br>(M = 1) |
| No. of TC<sup>b</sup> | 4–7<br>(M = 5) | 3–6<br>(M = 5) | 4–5<br>(M = 5) |
| No. of dorsal<br>kineties | 3 | 3 | 3 |
| No. of Ma<sup>b</sup> | 41–83<br>(M = 57) | 68–144<br>(M = 97) | 58–87<br>(M = 69) |
| Cortical<br>granules<sup>a</sup> | 0.5–1 µm, yellowish | 1–2 µm, yellowish | 0.7 µm; yellowish |
| Salinity (‰) | 10 | 4.3 | 4.1 |
| Data source | This work | Chen et al. (2013) | Jo et al. (2015) |

AZM, adoral zone of membranelles; BC, buccal cirri; FC, frontal cirri; FTC, frontoterminal cirri; LMR, left marginal row; M, mean; Ma, macronucleus; MVP, midventral pair; MVR, midventral row; No., number; PBC, parabuccal cirri; PTC, pretransverse cirri; RMR, right marginal row; TC, transverse cirri.

### Morphological comparison of the Italian population of Bakuella subtropica with the Chinese and Korean populations

The Italian population of *B*. *subtropica* closely resembles the Chinese type population, pop 1, and the Korean population (formerly *B*. *incheonensis*) in the ciliature pattern and other unique characteristics, that is having three frontal cirri, one buccal cirrus, one parabuccal cirrus, more than two frontoterminal cirri, dozens of midventral pairs, one or two midventral rows composed of three to five cirri, usually one pretransverse cirrus and five transverse cirri, three dorsal kineties, and the presence of yellowish cortical granules (Table 3; Chen et al., 2013; Jo et al., 2015). Furthermore, all these three populations were found in brackish water. However, there are some minor differences among them: (1) in terms of the body size of protargol stained specimens, the Chinese population is the largest (157 × 86 μm), the Italian population is middle-sized (138 × 50 μm), and the Korean population is the smallest (82 × 37 μm); (2) the number of adoral zone membranelles, frontoterminal cirri, midventral pairs, left marginal cirri, and right marginal cirri of the Italian specimens of *B*. *subtropica* are similar to those of the Chinese specimens, whereas all of these values are lower in comparison with the Korean specimens’ correspondent features; (3) the midventral complex of the Italian population occupies 76% of body length, which matches its length in the Chinese population (80%), but it is much shorter in the Korean population (62%); (4) the number of macronuclear nodules of the Italian population (41–83) matches well that of the Korean population (58–87), whereas the Chinese population possesses a much higher nodule number (on average, 97); (5) both the Italian and Korean populations possess similarly-sized cortical granules (about 0.7 μm), whereas the Chinese population shows 1–2 μm-sized cortical granules (Table 3). As all the above-mentioned characteristics have been proved to be highly variable between populations (Berger, 2006; Chen et al., 2020), we believe they are not significant for species-level discrimination. Thus, the identity of the studied organism is undoubtedly *B*. *subtropica*.

### Mitochondrial genome of the Italian population of Bakuella subtropica

As already reported in most sequenced ciliate mitogenomes (de Graaf et al., 2009; Swart et al., 2012; Serra et al., 2020), in *B*. *subtropica* we observed split 16S rRNA gene (*rnl*) and split protein coding genes (namely *nadh1*, *nadh2*, *rpS3*). Unfortunately, only in the case of *nadh2*, we have been able to retrieve both parts of the gene whereas in the other three cases we retrieved only one fragment, namely *nadh1*_*a*, *rpS3*_*b* and *rnl*_*a*. We hypothesized that the *nadn1*_*b*, *rpS3*_*a* and *rnl*_*b* subunits could not be identified due to high divergence from known reference sequences. Additionally, we also observed the split of *ccmf* gene that has been recently proposed as a typical feature (possibly a synapomorphy) of Hypotrichia and Euplotia (Zhang et al., 2021).

Another common feature found in other spirotrichs’ mitochondrial genome and in *B*. *subtropica,* is the presence of an AT-rich tandemly repeated region that roughly divides the genome into two halves and seems to indicate the origin of transcription directed towards the two edges (de Graaf et al., 2009; Swart et al., 2012; Serra et al., 2020). Besides, the mitogenome synteny is also significant among the representative hypotrichs in Fig. 3, except for the distinct inversion of *rpS10* in *B*. *subtropica*.

### Molecular phylogenetics and phylogenomics of Bakuella subtropica

Both phylogenetic and phylogenomic results confirmed the membership of *B*. *subtropica* to Urostylida and showed a close relationship with *U*. *grandis* (Figs. 4, 5). So far, there are only three available mitochondrial genomes of urostylids, hence, any discussion based on this dataset is still premature.

Our phylogenetic tree confirmed the clustering together of all sequences previously ascribed to *B*. *subtropica* and *B*. *incheonensis* with strong support (96/0.99). Moreover, it is consistent with previous studies (Chen et al., 2013; Jo et al., 2015; Lv et al., 2015; Lyu et al., 2018; Chen et al., 2020; Li et al., 2021) reporting *Bakuella* as a non-monophyletic genus in 18S rRNA gene-based phylogenetic tree. Indeed, all available *Bakuella* sequences are distributed into three separate clades, named clade 1, 2, and 3 (Fig. 4), suggesting that they might possibly represent independent genera. In clade 1, *B*. *litoralis* clustered with *Neobakuella flava* (type species) and *Apobakuella fusca* (type species) with moderate support (70/1.00), forming a sister assembly to *Diaxonella* spp. that includes the type species of the genus, *D*. *trimarginata* (49/0.97); in clade 2, *B*. *subtropica* populations were nested in a clade with three *Anteholosticha* species with a low support value (56/0.72); in clade 3, *Anteholosticha antecirrata*, *Bakuella granulifera*, *Bakuella guangdongica*, *Neobakuella aenigmatica*, an unidentified *Metabakuella* sp., and *U*. *grandis* (type species) were gathered together with strong support (99/1.00), and clade 3 was sister to clade 1 + 2 (96/1.00). Additionally, some phylogenetic assignments of species within these clades were also robustly supported in concatenated rDNA trees (Lv et al., 2015; Lyu et al., 2018; Xu et al., 2021). However, as shown in Table 4, the molecular evidence of the genera and the families involved in the three clades largely conflicts with morphological classification proposed by Berger (2006). Although a detailed discussion of the various observed discrepancies certainly falls outside the aim of our study and deserves a focused investigation, we could mention that only in clade 2 these species share some relatively consistent morphological features (Table 4). Indeed, *B*. *subtropica* is similar to *A*. *manca*, *A*. *multicirrata*, and *A*. *paramanca*, given that (1) some specimens of *B. subtropica* possess only one short midventral row composed of three cirri (Chen et al., 2013; present work) and interestingly, in *A. manca* sometimes this short midventral row (i.e., an atypical midventral pair composed of three cirri) may also be present (Berger, 2006); (2) all four species have only one buccal cirrus which may support the exclusion of *B*. *litoralis* (another species similar to *B*. *subtropica* in having few and short midventral rows, but more than one buccal cirrus) from clade 2 in the 18S rRNA gene phylogeny. This funding suggests that all four species in the clade 2 might possibly share a most recent common ancestor. However, since the molecular data of *Bakuella* species are rather insufficient, especially considering the absence of the type species (*Bakuella marina*) sequence, and that *Anteholosticha* is a well-known problematic group (Park et al., 2013; Fan et al., 2014; Fan et al., 2016; Lyu et al., 2018; Xu et al., 2021) with sequences scattered all over the urostylids, we consider premature any systematic reorganization of the involved species.

**Table 4.**
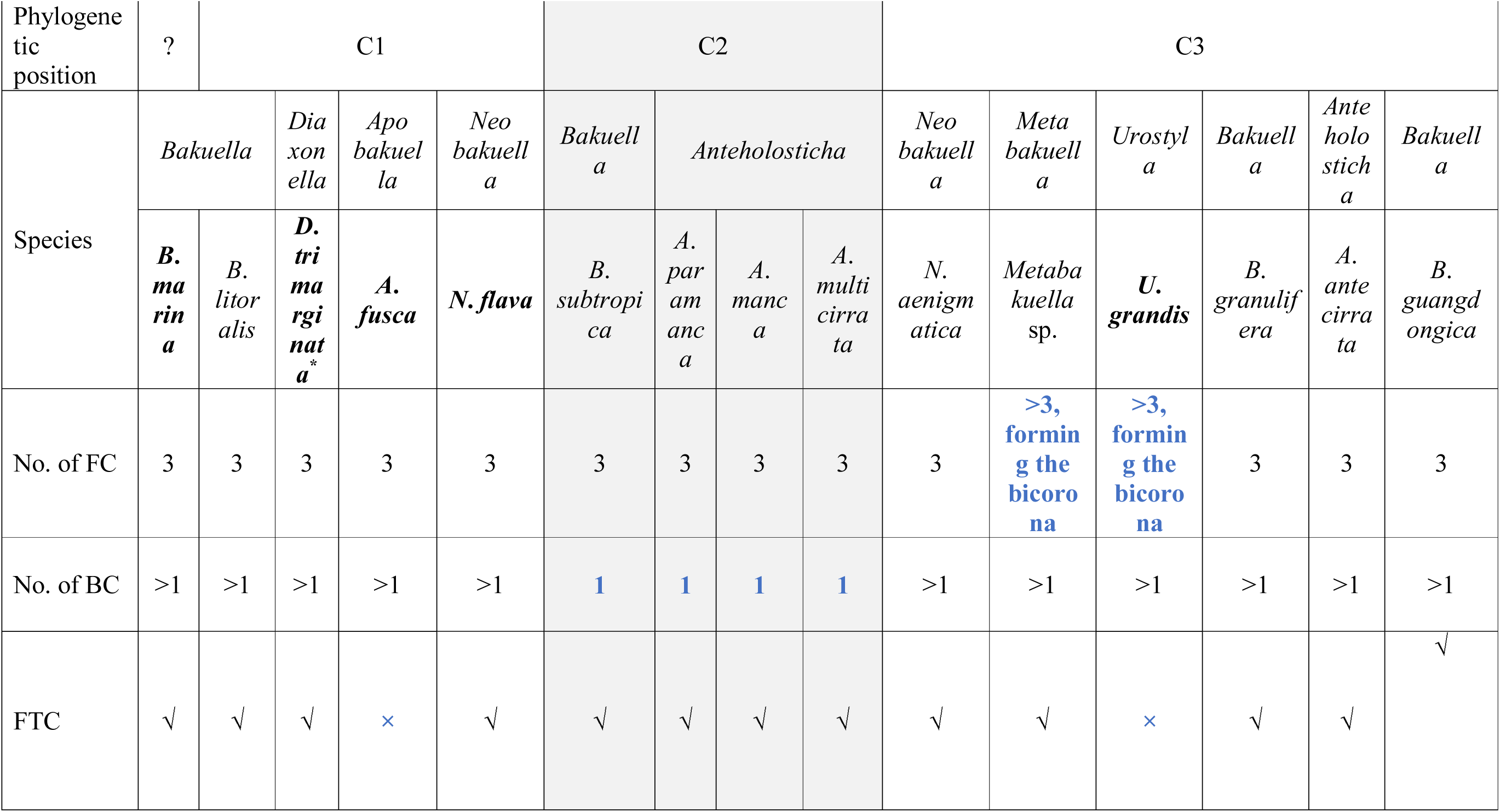
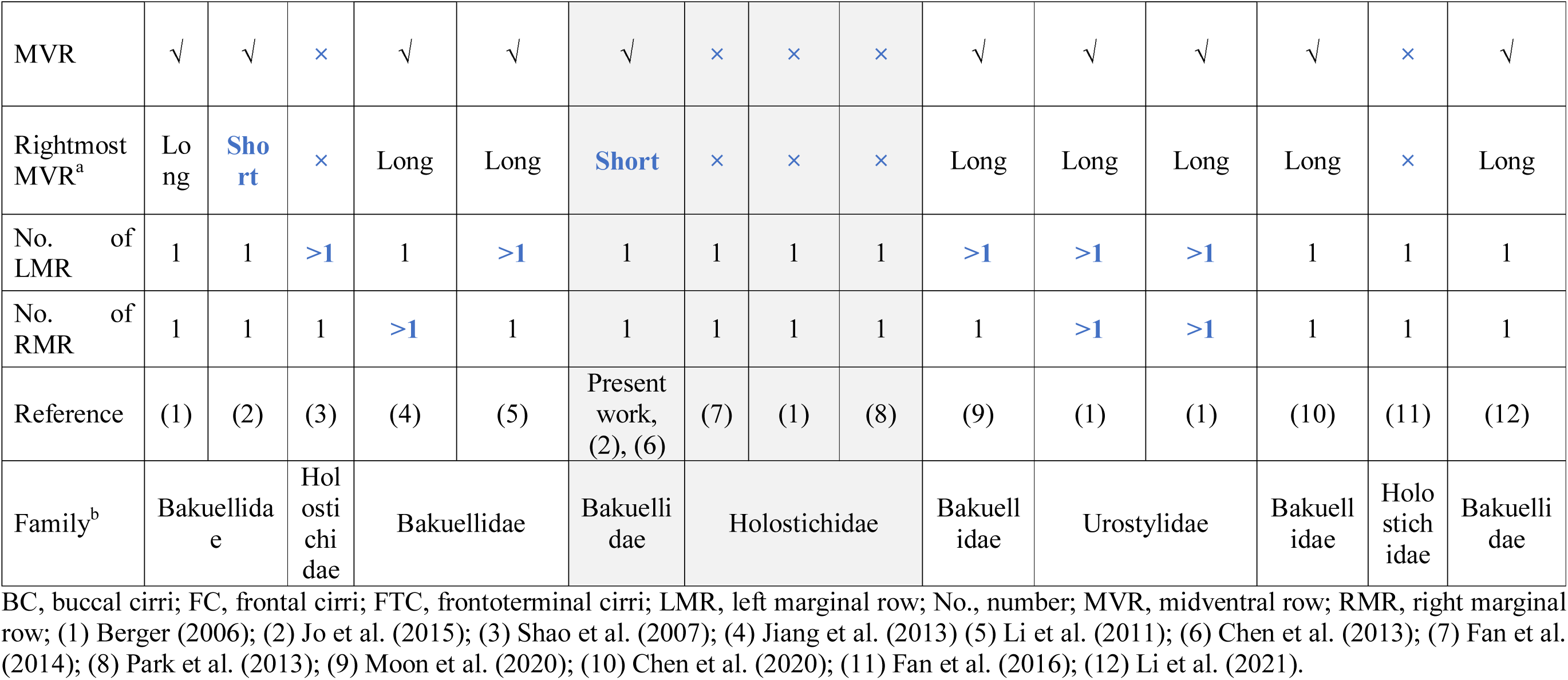
Comparison of related species/genera constituting the three clades (C1, C2, and C3) in the phylogenetic tree (Fig. 4) based on some selected main morphological characters. Species in C2, where *Bakuella subtropica* locates, are highlighted in grey shadow. Blue bold fonts highlight the morphological differences. Species in bold fonts are type species of the genus. ?, unknown; /, data unavailable; √, present; ×, absent; a, the short rightmost midventral row usually composed of three to five cirri, while the long one composed of more than five cirri; b, family classification based on Berger (2006); *, as for the four *Diaxonella* spp. in C1, only the type species, *D*. *trimarginata*, is listed.

To sum it up, as a member of the most confused groups of urostylids, the systematics of *Bakuella* and its relatives are still unresolved. This may be due to the lack of some important morphogenetic data and the convergent evolution of some diagnostic morphological features (Berger, 2006). An increased taxon sampling along with the acquisition of sequence data for additional gene markers, in combination with further understanding of the evolution of some main morphological characters, hopefully will help to solve these issues in the future.

### Comments on the phylogenetic position of Pseudourostyla cristata

*P*. *cristata*, the type species of *Pseudourostyla* Borror, 1972, shows the typical urostylid combination of morphological characters (Berger, 2006). Its systematic position inside the Urostylida core is robustly stable in all phylogenetic works based on nuclear genes (Lyu et al., 2018; Xu et al., 2021). However, in our phylogenomic work using mitochondrial genes, the position of *P*. *cristata* firstly breaks the previous rule, showing a close relationship to Stichotrichida, which is coherent with the latest work by Zhang et al. (2021) where, in total, 34 available ciliate mitogenomes were used. To solve the issue of the positioning of *P*. *cristata*, topology tests were performed to assess whether mitochondrial phylogenomic topology could be acceptable by 18S rDNA sequence-based tree and vice versa (Figs. S3, S4). Results confirmed that both the topologies retrieved from 18S rRNA gene dataset and mitochondrial dataset are statistically supported although they are not consistent with each other. In all cases topology tests rejected an alternative placement of *P*. *cristata*. The current inconsistency in the position of *P*. *cristata* revealed by nuclear and mitochondrial genes is most likely due to the paucity of available mitogenomes for some key urostylid representatives, although other hypotheses cannot be ruled out (e.g., it is worth remembering that sequences from the different markers have been produced in different laboratories using different populations of the organism). Only the availability of additional mitogenomes of urostylids and of *P*. *cristata* populations/strains will hopefully allow to solve the issue.

## Experimental procedures

### Description of the WWTP

A full-scale activated sludge WWTP managed and operated by Consortium Cuoiodepur S.p.A. (San Romano, Pisa, Italy) applying MLE process was selected in this study. The plant generally treats both domestic (3200 m^3^/d) and industrial (5900 m^3^/d) wastewater produced in the local tannery district. The biological section of this plant is constituted by an anoxic reactor (predenitrification, 11000 m^3^) and seven parallel aerobic reactors (nitrification, total 26000 m^3^) with an internal recirculation of ten times the influent flow rate (Figs. S5, S6) for removing nitrogen and organic compounds. Some operating parameters like HRT, SRT and volatile suspended solids concentration in the plant biological section are respectively 4 d, 70 d, and 8000 mg/L. HRT is about 2.8 d and 1.2 d in the nitrification and denitrification tank, respectively. From a biological point of view, this data mean that the microbial community is cyclically retained in average about 7 h in the nitrification tank and 3 h in the denitrification tank. Besides, the dissolved oxygen is controlled in the aerobic tanks at maintained around 2–3 mg/L.

### Sampling and wastewater characterizations

Samples of mixed liquors were periodically collected from the biological tanks in Cuoiodepur WWTP six times from June 15, 2018 until September 19, 2019. In detail, we routinely collected one bottle each time from the same points at the outlet of the aeration and anoxic reactors, and a total of 12 bottles of samples were collected. Samples were usually collected at mid-morning (around 10 a.m.), using plastic 1L-bottles. To meet the oxygen demand of samples collected from aeration tank, less than two-thirds of the volume of each bottle was occupied with the mixed liquor, and the bottles were left un-tightly lidded; on the contrary, samples taken from denitrification tank were fully filled and tightly lidded to ensure the permanence of the anoxic conditions. Samples were then transferred to laboratory within 3 h and three replicate subsamples were obtained from each original sample by stirring and pouring about 10 ml of liquid into 55 × 14 mm petri dishes (SARSTEDT, Nümbrecht, Germany). The subsamples were stood for about 15 min to settle the sludge particles; then microscopic analysis was performed within 12 h from the collection time (Dubber and Gray, 2009).

Meanwhile, throughout the sampling period, some abiotic parameters of the wastewater were monitored by the plant engineers via *in situ* detectors or following the APHA et al. (2012). From 2018 to 2019, the studied WWTP treated 1,437,430 m^3^ of industrial wastewater and 1,532,330 m^3^ of domestic wastewater in total per year. Table S2 reports the average values of the main physico-chemical parameters of three different water flows (namely, the industrial influent, the domestic influent and the final effluent discharged into the receiving water body) in Cuoiodepur throughout 2018, which were also similar in 2019. In addition, in Table S3, we showed some average physico-chemical parameters of the samples from the biological tanks one week before sampling. In terms of the 12 samples taken from nitrification and denitrification, their abiotic differences laid in (1) pH, since denitrification produces alkalinity, while nitrification consumes alkalinity; (2) soluble chemical oxygen demand, which is further removed in the aerobic reactor subsequently to the anoxic one; (3) there are some nitrate (about 10 mg/L) and no ammonium in the nitrification tank, while in the denitrification tank, the concentration of ammonium was higher, about 10–20 mg/L. Besides, the pH of the 12 samples was in general between 7–8, and the salinity was in the range of 5–10‰, which were measured respectively using the universal indicator paper (LLG-Labware, Meckenheim, Germany) and a refractometer (REICHERT-JUNG, New York, USA) after arrival at lab.

### Preliminary identification and enumeration of ciliates

Each subsample was firstly examined under a stereomicroscope (Leica MS5, Germany) at 10–40 × magnification, to observe the presence of living microorganisms and to check for the possible presence of small metazoans such as rotifers, nematodes, and other protists (except ciliates) such as flagellates and amoebae; these organisms were only registered without a precise identification as they were not a target of the present study. On the contrary, each living observed ciliate cell was isolated with a micropipette and moved from the petri dish to a glass slide to be examined under a light microscope (Leitz Weitzlar, Germany) equipped with a dedicated digital camera (Canon Power Shot S45), for preliminary identification using bright-field and differential interference contrast at 100–1,000 × magnification. As a reference, we used several taxonomic manuals, such as Curds (1982), Foissner and Berger (1996), and Kreutz and Foissner (2006) for species identification at this step when most observed species could be identified at least to family or even genus level. As for some taxa morphologically indistinguishable *in vivo* such as *Euplotes* spp., hypotrichs, and other doubtful species (i.e., those less familiar to us), these were preliminary operatively recorded with their known genus/family name + sp., or “Unknown hypotrich” + sp., or “Unknown ciliate” + sp., followed by a number (e.g., Unknown hypotrich sp. 1, Unknown hypotrich sp. 2, and so on), pending a further deeper investigation.

After the preliminary identification, within 12 h from sampling, we used a simple counting method to roughly (semi-quantitatively) estimate the relative abundance of each observed species. Specifically, we evaluated the relative abundance of each observed species based on the number of cells that could be collected in three subsamples within 30 min under the stereomicroscope. Finally, we defined four ciliate abundance categories: (1) highly abundant (i.e., dominant) species, when more than 30 cells could be collected in 30 min (we arbitrarily attributed to this category the highest score “3”); (2) abundant species, when 10 to 29 cells could be collected in 30 min (we arbitrarily attributed to this category the medium score “2”); (3) rare species, when less than 10 cells could be collected in 30 min (we arbitrarily attributed to this category the lowest score “1”); and (4) absent species, when no cell could be collected in 30 min (we arbitrarily attributed to this category the lowest score “0”). Apart from the aforementioned procedures at the sampling day, we also did a periodical check on non-clonal cultures maintained under laboratory culturing conditions in order to record the late-emerging species, regardless of their relative abundance, and this monitoring usually lasted two months. Finally, after sampling and monitoring periods (approximately from June 2018 to November 2019), the occurrence of all observed species (on the collection day and after) were summarized based on a presence-absence basis, together with information about the samples where they were found.

Moreover, in order to better understand the interrelationships between the species in the ciliate community and figure out the dynamic change on ciliate community during the sampling process, all observed species were (1) divided according to their habits into three functional groups: (1.1) free-swimming forms, which swim actively throughout the liquid phase of the mixed liquor; (1.2) crawling forms, which crawl over the surface of the sludge flocs; (1.3) sessile forms, which adhere, usually by means of a stalk, to the sludge flocs (Curds, 1973a); and (2) annotated with their types of feeding behaviour into the following categories: (2.1) filter feeding ciliates, obligatory bacterivores (such as many scuticociliates, peritrichs); (2.2) carnivorous ciliates, true predators (such as haptorians, suctorians, which are obligatory feeding on other ciliates, or even small metazoans); (2.3) omnivorous ciliates (which are also filter-feeders but, in addition to bacteria, they can also engulf/feed small ciliates, or organic detrital pieces). These trophic categories were slightly modified respect to those proposed by previous papers (e.g., Madoni, 1994; Foissner and Berger, 1996; Rosati et al., 2008).

### Ciliate culturing and sample maintenance

Rice grains were added to the original samples and subsamples after microscopic analysis to support the growth of indigenous bacteria and flagellates as food source for the existing ciliate species and stimulating the excystment of other, possibly present, encysted species (Foissner, 1992). Such non-clonal cultures were monitored according to the following frequencies: one week, two weeks, one month, and two months after the first culture enrichment, until no additional new species appeared. Meanwhile, we tried to establish monoclonal or polyclonal cultures of all observed species useful for further analyses. Considering the average habitat salinity, cultivations were performed in 10‰ salinity medium by diluting artificial seawater (salinity about 33‰, produced dissolving the Red Sea Salt (Red Sea Europe, Verneuil-sur-Avre, France) with mineral water (Fonte Monteverde, Pracchia, Italy) following manufacturer’s protocols). The monoclonal cultures of the green flagellate *Dunaliella tertiolecta* (original culturing salinity 5‰) and of *Raoultella planticola* (*Gammaproteobacteria*) (original culturing salinity 0‰, cultivated in Cerophyll medium; see Boscaro et al. (2012) for details) were brought to 10‰ salinity using the salinity medium, and then added to the cultures as a food resource. However, the polyclonal cultures of the suctorian *P*. *libera* were fed with monoclonal cultures of *P*. *calkinsi*. All cultures were fed once a week. All original samples and cultures were maintained in incubator at a temperature of 19 ± 1°C over the entire period of investigation.

### Follow-up morphological analysis of ciliates

To morphologically characterize and identify the ciliates, appropriate staining techniques like silver impregnation (protargol) (Wilbert, 1975), silver carbonate method (Fernández-Galiano, 1976), wet silver nitrate method (Chatton and Lwoff, 1930), and Feulgen nuclear reaction (Dragesco and Dragesco-Kernèis, 1986) were selectively performed depending on the species and based on Foissner (2014), for studying the ciliature and the silverline pattern; these techniques were also combined with molecular techniques (see below). For these studies, ciliates were collected either from non-clonal samples or from the later successfully established mono/polyclonal cultures depending on their cultivability.

Optical microscopy pictures were captured at 100–1,000 × magnification using a light microscope (Leitz Weitzlar, Germany) equipped with a digital camera (Canon Power Shot S45); images were used to obtain cell dimensions of living and stained ciliates. Morphometric measurements were analyzed with ImageJ 1.46r software (Ferreira and Rasband, 2012). CombineZM and Adobe Photoshop CS6 software were used for photograph processing.

### Follow-up molecular analysis of ciliates

For abundant or cultivable species, standard DNA extraction was performed using the NucleoSpin™ Plant II DNA extraction kit (Macherey-Nagel GmbH and Co., Düren NRW, Germany). A reasonable number (usually 30–70) of conspecific cells collected either from non-culturable subsamples (according to preliminary identification) or later established cultures were isolated and manipulated using a glass micropipette, washed twice in 10‰ salinity medium and once in distilled water, and then fixed in 70% (v/v) ethanol at −20°C for DNA extraction following the manufacturer’s instructions. For nonculturable ciliate species or species to be processed through downstream analyses (e.g., next generation sequencing), the REPLI-g Single Cell Kit (QIAGEN®, Hilden, Germany) was used, starting from few (1–30) conspecific cells isolated either from non-culturable subsamples or later established cultures.

As for the Italian population of *B*. *subtropica*, a single cell was isolated from the laboratory culture, and then washed three times before putting it into PBS buffer. Then, the cell was transferred into a 0.2 ml tube (Eppendorf, Hamburg, Germany) together with 4 μl of PBS. The whole genome amplification (WGA) protocol was completed following the REPLI-g Single Cell Kit protocol, according to manufacturer’s instructions.

Polymerase chain reaction (PCR) was performed in a C1000™ Thermal Cycler (Bio-Rad, Hercules, USA). The almost full-length of the 18S rRNA gene was amplified using the primers 18S F9 (Medlin et al., 1988) and 18S R1513 Hypo (Petroni et al., 2002). High-fidelity Takara Ex Taq PCR reagents were employed (Takara Bio Inc., Otsu, Japan) according to the manufacturer’s instructions. PCR cycles were set as follows: 3 min 94°C, 35 × [30 s 94°C, 30 s 55°C, 2 min 72°C], 6 min 72°C. PCR products were purified with the EUROGOLD Cycle-Pure Kit (EuroClone, Milan, Italy) and subsequently sent for direct Sanger sequencing to an external sequencing company (GATC Biotech AG, European Custom Sequencing Centre, Germany) by adding appropriate internal primers 18S R536, 18S F783, and 18S R1052 (Nitla et al., 2019). Additionally, the ITS and the 28S rRNA gene sequencing of *B*. *subtropica* were obtained according to Liao et al. (2020).

### Next generation sequencing, mitochondrial genome assembly and annotation of Bakuella subtropica

After verifying the correctness and quality of the WGA material of *B*. *subtropica* Italian population (see above) via 18S rRNA gene amplification and later Sanger sequencing, this material was processed with a Nextera XT library and sequenced at Admera Health (South Plainfield, USA), using Illumina HiSeq X technology to generate 63,387,611 reads (paired-ends 2 × 150 bp). Preliminary assembly of resulting reads was performed using SPAdes software (v 3.6.0) (Bankevich et al., 2012). Contigs representing mitochondrial genome were identified using the Blobology pipeline (Kumar et al., 2013), and by TBLASTN searches using as queries proteins from reference genomes downloaded from NCBI, namely *P*. *cristata* (MH888186) and *Laurentiella strenua* (KX529838). Contigs with a GC content comprised between 25% and 35%, and a coverage higher than 1000 × were selected and a subset of the extracted reads was assembled with SPAdes. We decided to use approximately 10% of the extracted reads to artificially reduce the reads coverage, as coverage above 100 × generally produces worse assemblies. Indeed, a too high coverage may cause an increasing number of exactly replicated sequencing errors, which create false branches in the de Bruijn graphs. The assembled genome was annotated using Prokka 1.10 (Seemann, 2014), setting the DNA translation codon table “4” and then manually checked.

### Phylogenetic and phylogenomic analyses on Bakuella subtropica

The 18S rRNA gene sequence of *B*. *subtropica* was assembled using Chromas Lite 2.1 software and compared with the non-redundant sequence database using NCBI-BLAST; then aligned with the automatic aligner of ARB software package version 5.5 (Westram et al., 2011) together with related sequences contained in the SSU ref NR99 SILVA database (Quast et al., 2013) and some relevant latest released sequences on GenBank. For the phylogenetic analyses based on18S rRNA gene, a total of 126 sequences was selected. In addition to *B*. *subtropica* Italian population, 119 sequences belonging to the subclass Hypotrichia were selected as ingroup, plus four choreotrichs and two oligotrichs as outgroup. All selected 18S rRNA sequences were aligned again using the automatic aligner of ARB package version 5.5 and then manually checked. After trimming at the shortest sequencing length, a positional filter was applied to retain those columns with at least a 10% of similarity, and at the end, producing a 1,577-character matrix. The GTR + I + G was selected as best model for the phylogenetic analyses using jModelTest 2 (Darriba et al., 2012). ML tree was inferred using PHYML software version 2.4 (Guindon and Gascuel, 2003) from the ARB package, setting 1,000 pseudo replicates for statistical support. BI tree was instead inferred with MrBayes version 3.2 (Ronquist et al., 2012), using the same substitution model as in the ML analysis. Three different Markov chain Monte Carlo runs were used, with one cold chain and three heated chains, with a burn-in of 25%, iterating for 5,000,000 generations to reach an appropriate confidence level.

For the phylogenomic analysis of *B*. *subtropica* based on the complete mitochondrial genome, other 11 available spirotricheans’ sequences on GenBank have been downloaded and added as ingroup; five oligohymenophorean sequences were selected as outgroup, among them, two *Paramecium* mitochondrial genomes (namely *Paramecium biaurelia* strain V1-4 and *Paramecium caudatum* strain 43c3d) were additionally downloaded from ParameciumDB (Arnaiz et al., 2020). Specifically, 18 protein coding genes with at least 80% presence in the mitochondrial genomes of the above mentioned 17 species were selected for the analysis. Amino acidic sequences for each set of genes were aligned using MAFFT (Katoh and Standley, 2013). Then, we concatenated the multi-alignments of all 18 genes (gaps were inserted in few rare cases where a protein was missing from a certain species), and finally obtained the employed matrix composed by 11,587 sites. The CAT substitution model, with MtArt as replacement matrix, was estimated as the best with ProtTest version 3.2 (Darriba et al., 2011). RaxML software version 8 (Stamatakis, 2014) was used to reconstruct the ML phylogenetic tree using 100 bootstraps.

### Terminology and systematics of ciliates

Systematic classification for the phylum Ciliophora is mainly based on Adl et al. (2019). For the subclass Hypotrichia, terminology used is according to Berger (2006), and systematics is based on both Adl et al. (2019) and Xu et al. (2021).

## Acknowledgements

This work was supported by the European Commission H2020-PEOPLE-RISE project NGTax (872767) (to G.P.); the University of Pisa PRA_2018_63 project (to G.P.); the Ministry of Foreign Affairs and International Cooperation Scholarships Protocol ID:1559731487 (to W.L.).

Great thanks to: Consorzio Cuoiodepur, for providing the sampling materials; Network Inter-Library Document Exchange system, and, especially, Mrs. Barbara Lapucci, the staff of the Biblioteca di Scienze Naturali e Ambientali di Pisa, for providing lots of old literature references; Mr. Alessandro Allievi for his kind help in drawing the illustration (Fig. 6); and Dr. Venkatamahesh Nitla for his help in obtaining the sequence of *Metopus*.

## Author Contributions

G.P., G.Mu., G.Mo, and L.M. conceived the study; G.P., L.M., L.G., V.S., and W.L. designed the experiments; F.S., L.G., and W.L. performed the sampling; F.S. and G.Mo. monitored the physico-chemical parameters of the studied WWTP; W.L. cultured, identified, and semi-quantified the ciliates present in samples; L.G. and W.L. undertook the molecular experiments and analyses, and, specifically, L.G. undertook the mitochondrial genomic analysis; G.P. and L.M. supervised the research; G.P. financed the research; W.L. wrote the original draft and all co-authors contributed with writing and/or with critical feedback to the final version.

## Supporting Information

Additional Supporting Information may be found in the online version of this article at the publisher’s web-site: XXXXXXX.

