## Supplemental Table 1-3 and Supplemental Figure 1-6 for "Ciliated protozoa from industrial WWTP activated sludge: a biodiversity survey including trophic interactions and redescription of *Bakuella subtropica* (Spirotrichea, Hypotrichia) according to Next Generation Taxonomy"

**Table S1.** List of species observed in Cuoiodepur WWTP with their relative abundance scores at the different sampling days and different tanks. The species collected on the sampling day are displayed in the upper part of the table, then the species appeared only later during lab maintenance (shadowed in grey) are listed. Species order refers to Table 1 in the main text. The relative abundance scores of the species observed are presented as follows: 0 = absence in three subsamples on the sampling day; 1= present in few numbers (i.e., 1 to 9 cells could be collected from three subsamples within 30 min); 2 = present in moderate numbers (i.e., 10 to 29 cells could be collected within 30 min); 3 = present in massive numbers (i.e., more than 30 cells could be collected within 30 min). Asterisks indicate the original samples where the species of interest appeared after laboratory enrichment. Note that, since the observed species of *Euplotes* were not discriminable *in vivo* due to their high morphological similarity, they were grouped as a single taxonomical unit (*Euplotes* spp.). In detail, the four observed *Euplotes* species were *Euplotes* sp. 1, *Euplotes* sp. 2, *E*. *curdsi*, and *E*. *vanleeuwenhoeki*, and they were observed throughout the study, with their first record on 15-Jun-2018, 15-Jun-2018, 4-Jul-2019, and 19-Sep-2019, respectively.

| Species name (average size) | | 15-Jun-18  (Ⅰ) | | 7-Dec-18  (ⅠⅠ) | | 15-Mar-19  (ⅠⅠⅠ) | | 18-Apr-19  (ⅠⅤ) | | 4-Jul-19  (Ⅴ) | | 19-Sep-19  (ⅤⅠ) | |
| --- | --- | --- | --- | --- | --- | --- | --- | --- | --- | --- | --- | --- | --- |
|  |  | Ni | De | Ni | De | Ni | De | Ni | De | Ni | De | Ni | De |
| *Aspidisca* cf.  *cicada* (25 × 20 μm) | | 1 | 0 | 1 | 1 | 1 | 1 | 2 | 2 | 0 | 1 | 1 | 1 |
| *Euplotes* spp. | *Euplotes* sp. 1 (60 × 30 μm) | 1 | 0 | 0 | 0 | 1 | 0 | 2 | 2 | 3 | 3 | 3 | 3 |
|  | *Euplotes* sp. 2 (40 × 30 μm) |  |  |  |  |  |  |  |  |  |  |  |  |
|  | *E*. *curdsi* (65 × 40 μm) |  |  |  |  |  |  |  |  |  |  |  |  |
|  | *E*. *vanleeuwenhoeki* (55 × 30 μm) |  |  |  |  |  |  |  |  |  |  |  |  |
| *Bakuella* *subtropica* (135 × 40 μm) | | 1 | 0 | 0 | 0 | 0 | 0 | 1 | 1 | 0 | 0 | 1 | 0 |
| Unknown hypotrich sp. 1 (75 × 25 μm) | | 0 | 0 | 1 | 0 | 1 | 1 | 0 | 0 | 0 | 0 | 1 | 1 |
| Unknown hypotrich sp. 2 (55 × 10 μm) | | 0 | 0 | 1 | 0 | 2 | 1 | 2 | 2 | 3 | 1 | 0 | 0 |
| *Pseudochilodonopsis* cf. *mutabilis* (40 × 25 μm) | | 1 | 0 | 1 | 1 | 1 | 1 | 0 | 0 | 0 | 1 | 0 | 1 |
| *Cyclidium* cf. *marinum* (20 × 15 μm) | | 0 | 0 | 0 | 0 | 0 | 0 | 0 | 0 | 0 | 0 | 3 | 3 |
| *Paramecium calkinsi* (110 × 40 μm) | | 1 | 0 | 3 | 3 | 1 | 2 | 3 | 3 | 0 | 0 | 1 | 1 |
| *Holophrya teres* (120 × 70 μm) | | 3 | 3 | 0 | 0 | 3 | 3 | 1 | 1 | 1 | 1 | 0 | 0 |
| *Phialina* sp. (80 × 20 μm) | | 2 | 2 | 2 | 2 | 2 | 2 | 3 | 3 | 0 | 0 | 2 | 1 |
| *Trochiliopsis* *australis* (30 × 15 μm) | | 3 | 3 | 1 | 0 | 1 | 1 | 0 | 0 | 0 | 0 | 0 | 0 |
| *Pseudovorticella spathulata*  (37 × 27 μm, adult without a stalk) | | 0 | 1 | 1 | 0 | 0 | 0 | 0 | 0 | 2 | 3 | 3 | 2 |
| *Thuricola* *similis* (195 × 15 μm, zooid) | | 3 | 3 | 0 | 0 | 1 | 1 | 1 | 1 | 0 | 0 | 0 | 0 |
| *Acineria uncinata* (35 × 10 μm) | | 0 | 0 | 0 | 0 | 0 | 0 | * | * | 0 | 0 | 0 | 0 |
| *Zosterodasys* sp. (105 × 25 μm) | | * | * | 0 | 0 | * | * | * | * | 0 | 0 | 0 | 0 |
| *Metopus* sp. (140 × 40 μm) | | 0 | * | 0 | 0 | 0 | 0 | 0 | * | 0 | 0 | 0 | 0 |
| *Uronema* sp. (25 × 10 μm) | | 0 | 0 | 0 | 0 | * | * | * | * | * | * | 0 | 0 |
| *Podophrya libera* (adult without a stalk, about 25 μm across) | | 0 | 0 | 0 | 0 | 0 | * | 0 | 0 | 0 | 0 | 0 | 0 |

De, denitrification tank; Ni, nitrification tank; Ⅰ–ⅤⅠ, the first to sixth sampling date.

**Table S2.** Characterization of the main physico-chemical parameters of three different water streams in Cuoiodepur WWTP throughout 2018. Data are shown as mean value ± standard deviation.

| Source  Parameter | Industrial influent | Domestic influent | Effluent |
| --- | --- | --- | --- |
| pH | 8.4 ± 1.3 | 7.5 ± 0.5 | 7.6 ± 0.4 |
| COD total (mg/L) | 12,226 ± 3,974 | 245 ± 96 | 138 ± 19 |
| COD soluble (mg/L) | 5,705 ± 3,183 | / | / |
| TOC (mg/L) | 1,837 ± 493 | / | / |
| N-NH_4_^+^ (mg/L) | 293 ± 142 | 25 ± 14.8 | 2.2 ± 2.5 |
| N-NO_3_^-^ (mg/L) | / | / | 7.5 ± 8.2 |
| TKN (mg/L) | 809 ± 262 | 50.4 ± 12.4 | 19.4 ± 4.6 |
| TP (mg/L) | 22.9 ± 14.4 | 3.2 ± 1.1 | 0.3 ± 0.06 |
| HS^-^ (mg/L) | 253 ± 194 | / | / |
| S-SO_4_^2-^ (mg/L) | 2,304 ± 502 | 54 ± 32 | 1,389 ± 173 |
| Cl^-^ (mg/L) | 6,279 ± 1,852 | 218 ± 183 | 2,930 ± 548 |
| TSS (mg L^-1^) | / | / | 3.2 ± 1.9 |

/, data unavailable; COD, chemical oxygen demand; TKN, total Kjeldahl nitrogen; TOC, total organic carbon; TP, total phosphorous; TSS, total suspended solids.

**Table S3.** Characterization of the main physico-chemical parameters (average) of biological tanks in the week before each sampling time (I–VI).

| Parameter | Sampling date | | | | | |
| --- | --- | --- | --- | --- | --- | --- |
|  | 15-Jun-18  (I) | 7-Dec-18  (II) | 15-Mar-19 (III) | 18-Apr-19  (IV) | 4-Jul-19  (V) | 19-Sep-19  (VI) |
| Temperature (°C) | 31.5 | 26.2 | 26.0 | 26.8 | 36.6 | 33.3 |
| pH | 7.0 | 7.3 | 7.5 | 7.4 | 7.5 | 7.2 |
| Cl^-^ (mg/L) | 2,913.0 | 3,049.2 | 3,101.8 | 2,979.3 | 3,494.0 | 3,312.0 |
| S-SO_4_^2-^ (mg/L) | 1,548.8 | 1,664.0 | 1,585.6 | 1,565.5 | 1,723.7 | 1,621.5 |
| COD soluble (mg/L) | 529.6 | 532.8 | 536.0 | 587.0 | 609.3 | 760.0 |
| N-NH_4_^+^ (mg/L) | 4.2 | 4.5 | 3.9 | 3.7 | 6.1 | 4.3 |
| MLSS (mg/L) | 8,468 | 12,824 | 12,336 | 12,835 | 11,207 | 10,245 |
| MLVSS (mg/L) | 6,808 | 8,746 | 9,153 | 9,883 | 8,106 | 7,581 |

COD, chemical oxygen demand; MLSS, mixed liquor suspended solids; MLVSS, mixed liquor volatile suspended solids.

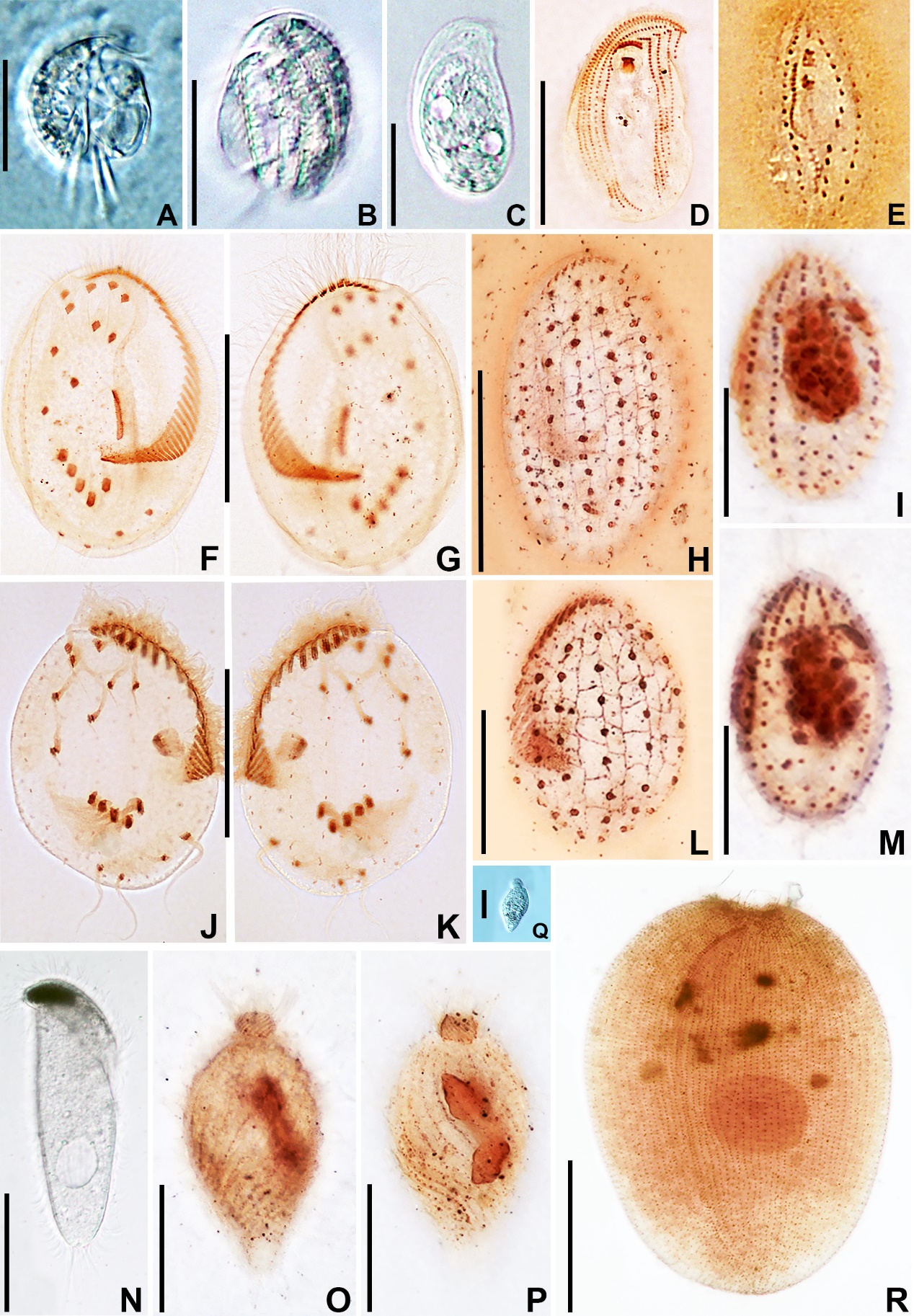

**Figure S1.** Photomicrographs of some species found in Cuoiodepur WWTP-part I. A, B. *Aspidisca* cf. *cicada in vivo*, showing the ventral (A) and dorsal characteristics (B). C, D. *Pseudochilodonopsis* cf. *mutabilis* *in vivo* (C, ventral side) and after protargol staining (D, ventral ciliature pattern). E, I, M. *Uronema* sp. (E, details of oral apparatus after silver nitrate staining; I, M, somatic kineties on both sides of the cell after protargol staining). F–H. *Euplotes* sp. 1 after protargol staining (F, ventral ciliature pattern and G, dorsal ciliature pattern), and after silver nitrate staining (H, silverline pattern on the dorsal side). J–L. *Euplotes* sp. 2 after protargol staining (J, ventral ciliature pattern and K, dorsal ciliature pattern), and after silver nitrate staining (L, silverline pattern on the dorsal side). N. *Metopus* sp. *in vivo*. O–Q. *Phialina* sp. after protargol staining (O, P, somatic kineties on both sides of the cell and its macronucleus) and *in vivo* (Q, a slightly contracted cell). R. *Holophrya teres* after silver carbonate staining, showing the ciliary rows, the three-rowed dexiotrop brosse, and the nuclear apparatus. Scale bars: 10 μm (I, M), 20 μm (A–D, L, O–Q), 40 μm (F–H, J, K), 60 μm (N, R).

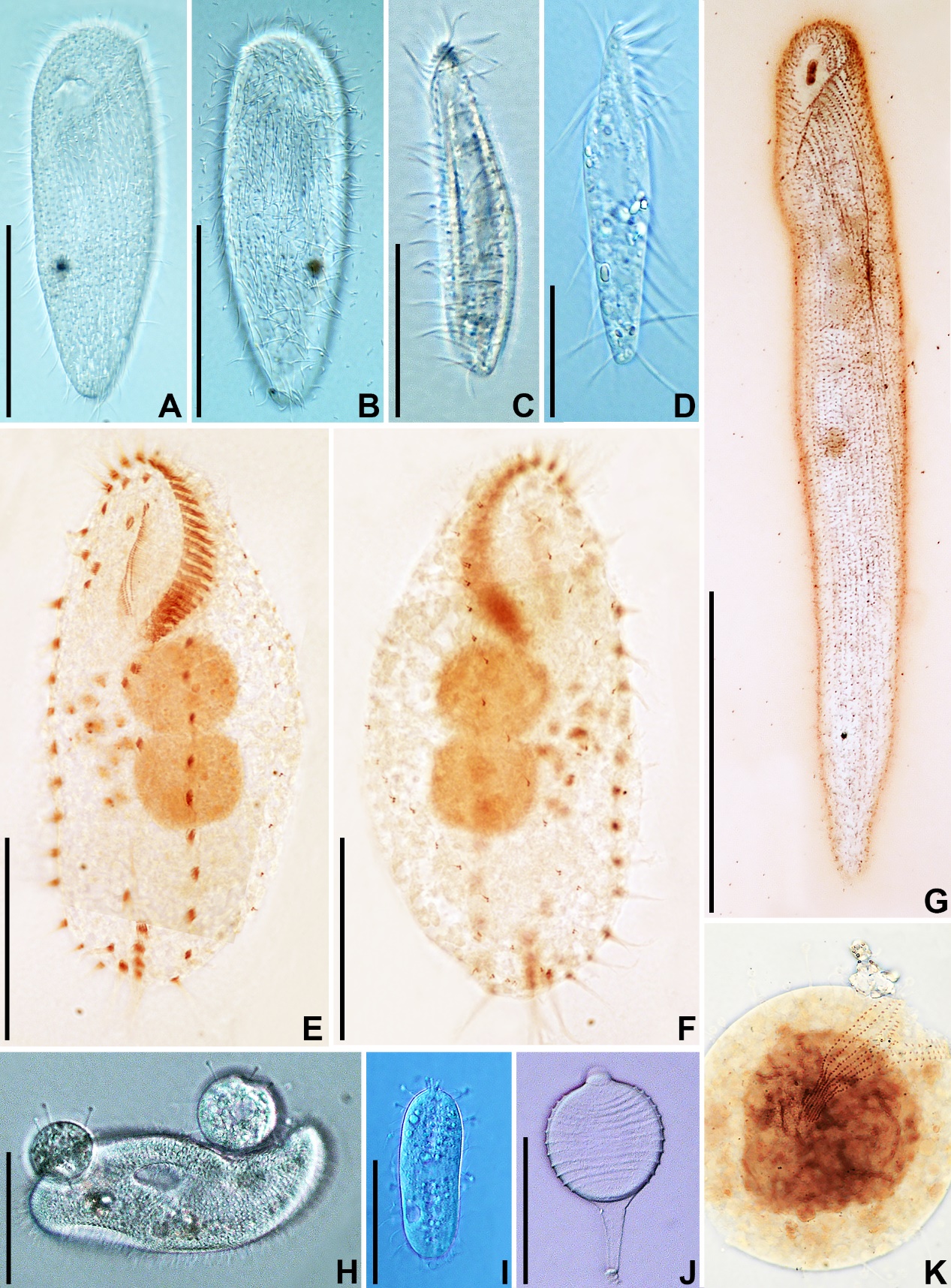

**Figure S2.** Photomicrographs of some species found in Cuoiodepur WWTP-part II. A, B, G. *Zosterodasys* sp. *in vivo* showing the ventral (A) and dorsal characteristics (B), and a rather long cell after silver nitrate staining (G, ventral ciliature pattern). C. *Acineria uncinata in vivo*. D. Unknown hypotrich sp. 2 *in vivo*. E, F. Unknown hypotrich sp. 1 after protargol staining showing ventral (E) and dorsal (F) ciliature patterns, as well as the nuclear apparatus. H–K. *Podophrya libera in vivo* (H–J) and after protargol staining (K) showing different stages during life cycle: adult (H, two spherical cells are preying on *Paramecium calkinsi*), swarmer (I), mature cyst (J), and an early divider (K). Scale bars: 20 μm (C–F); 40 μm (A, B, H–J); 80 μm (G).

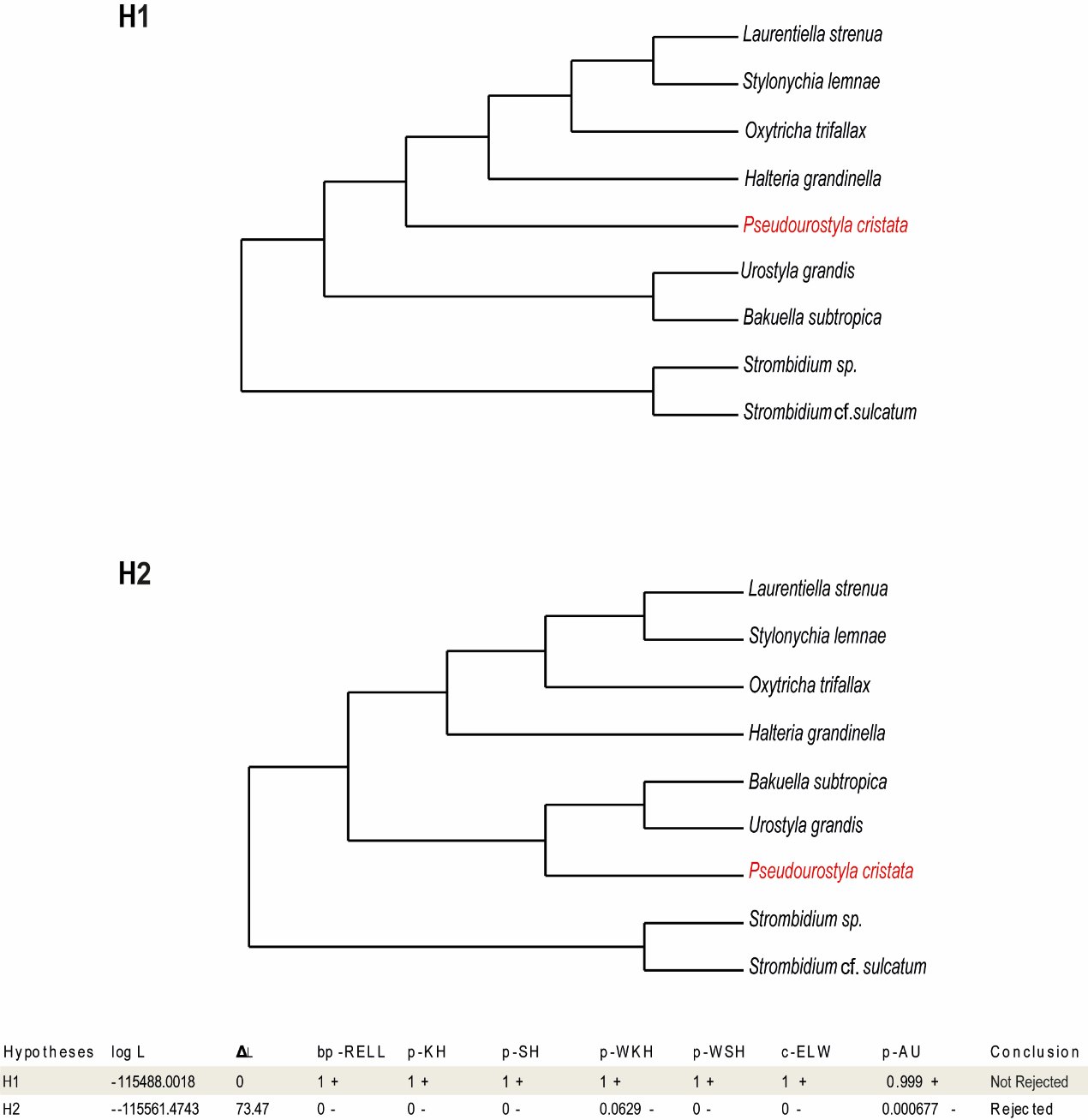

**Figure S3.** Alternative mitochondrial based phylogenomic hypotheses regarding different positions of *Pseudourostyla cristata* (highlighted in red font). Two hypotheses (H1 and H2) are presented, where H1 corresponds to the retrieved ML tree (Fig. 5) and H2 represents the topology that *P*. *cristata* is located inside Urostylida to mirror the result obtained from 18S rRNA gene analysis (Fig. 4). The topological test results of the two hypotheses are shown at the bottom where plus signs denote the 95% confidence set, and minus signs denote significant exclusion. logL, log-likelihood; ΔL, logL difference from the maximal logL in the set; bp-RELL: bootstrap proportion using RELL method; p-KH, p-value of one-sided Kishino-Hasegawa test; p-SH, p-value of Shimodaira-Hasegawa test; p-WKH, p-value of weighted KH test; p-WSH: p-value of weighted SH test; c-ELW, Expected Likelihood Weight; p-AU, p-value of approximately unbiased test. Topology tests were performed on the IQ-TREE software (version 1.6.12) (Nguyen et al., 2015) using RELL method (Kishino et al., 1990) with 10,000 replicates.

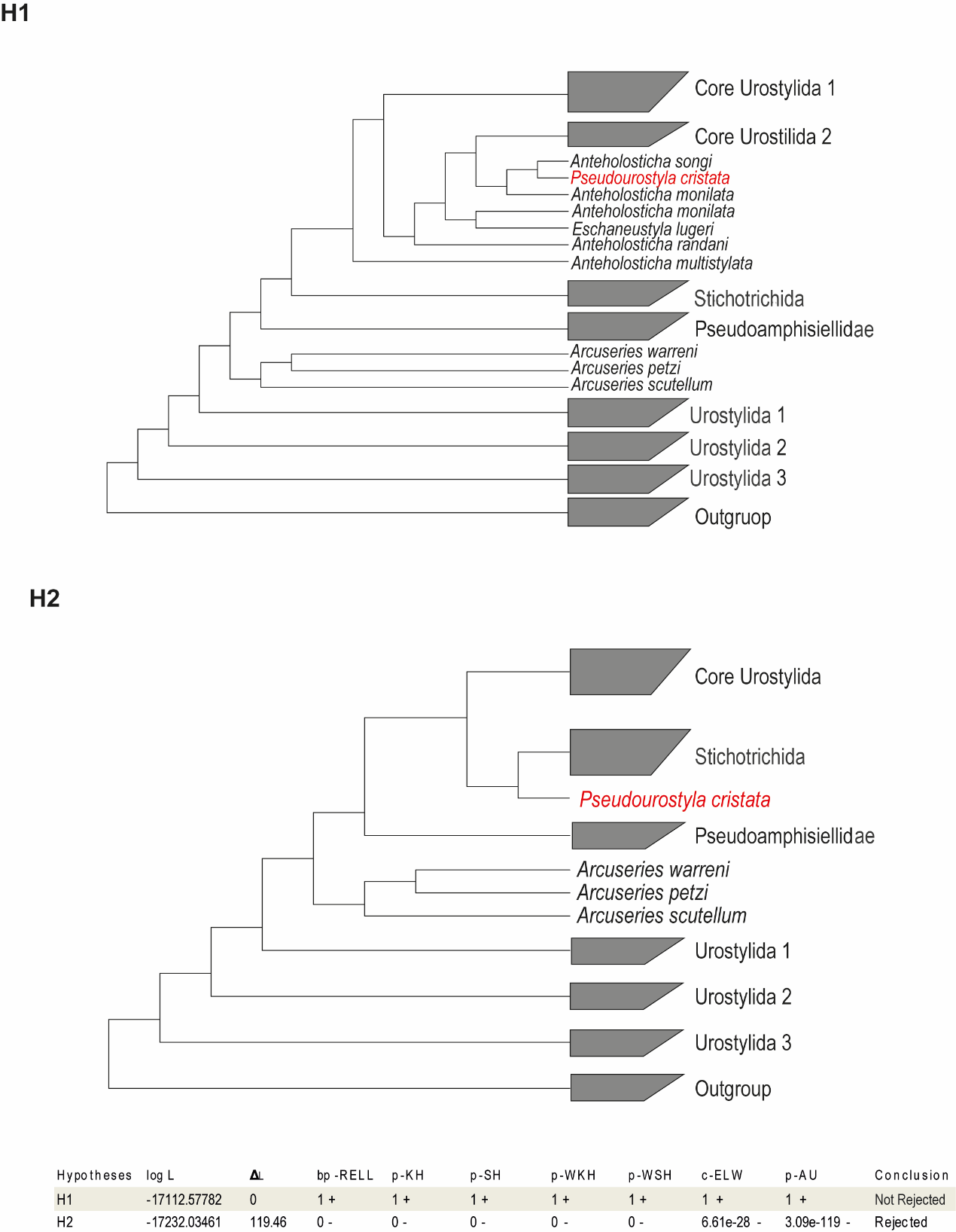

**Figure S4.** Alternative 18S rRNA based phylogenetic hypotheses regarding different positions of *Pseudourostyla cristata* (highlighted in red font). Two hypotheses (H1 and H2) are presented, where H1 corresponds to the retrieved ML/BI tree (Fig. 4) and H2 represents the topology that *P*. *cristata* is located as an early branch from Stichotrichida to mirror the result obtained from mitochondrial genome analysis (Fig. 5). The topological test results of the two hypotheses are shown at the bottom where plus signs denote the 95% confidence set, and minus signs denote significant exclusion. logL, log-likelihood; ΔL, logL difference from the maximal logL in the set; bp-RELL: bootstrap proportion using RELL method; p-KH, p-value of one-sided Kishino-Hasegawa test; p-SH, p-value of Shimodaira-Hasegawa test; p-WKH, p-value of weighted KH test; p-WSH: p-value of weighted SH test; c-ELW, Expected Likelihood Weight; p-AU, p-value of approximately unbiased test. Topology tests were performed on the IQ-TREE software (version 1.6.12) (Nguyen et al., 2015) using RELL method (Kishino et al., 1990) with 10,000 replicates.

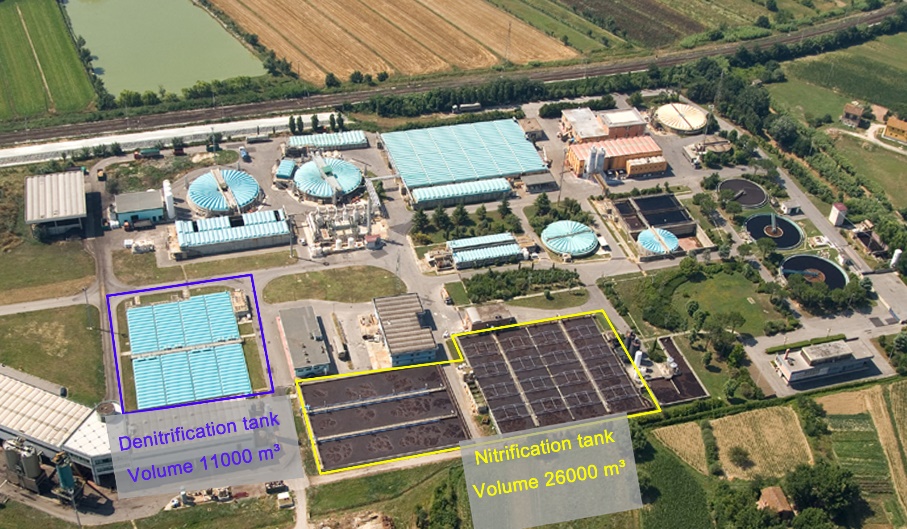

**Figure S5.** Cuoiodepur S.p.A. WWTP, with emphasis on the biological tanks.

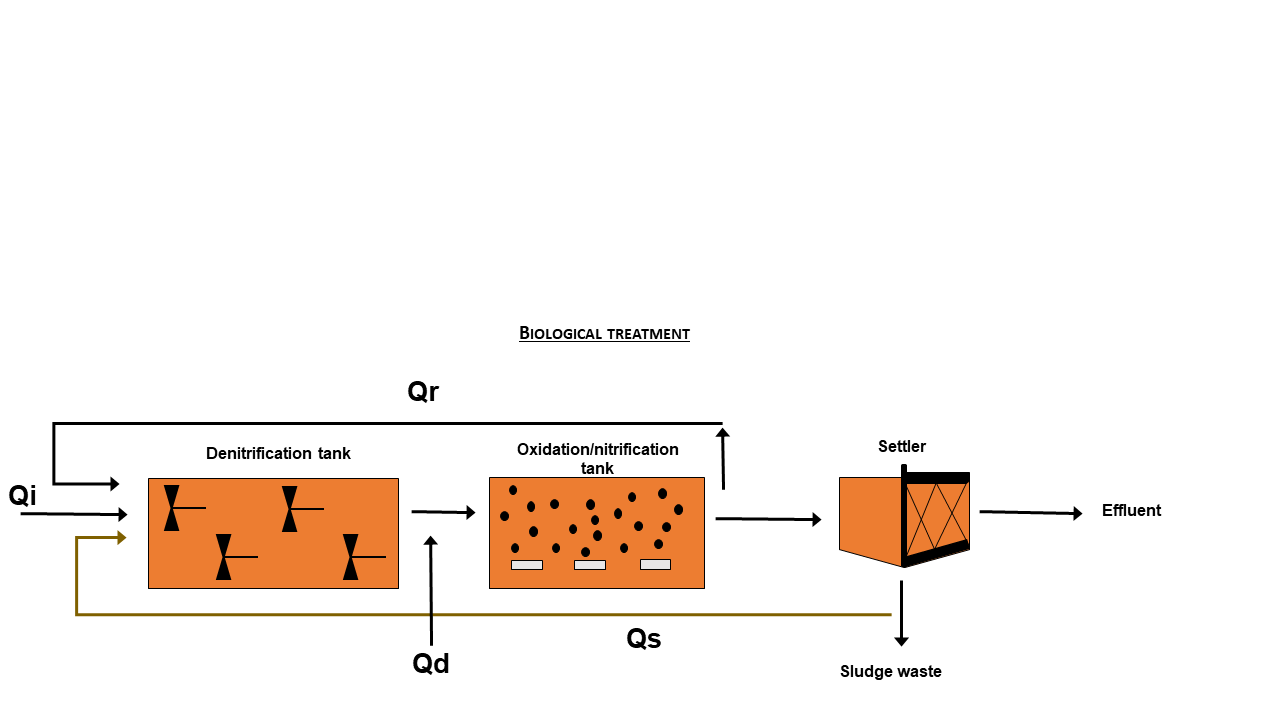

**Figure S6.** Biological treatment process in Cuoiodepur S.p.A. WWTP. Qd, flow of domestic wastewater; Qi, flow of industrial wastewater; Qr, flow of internal recirculation (10 times of Qi); Qs, flow of recycled activated sludge (1.5 times of Qi).
